# Longitudinal Quantification of Metabolites and Macromolecules Reveals Age- and Sex-Related Changes in the Healthy Fischer 344 Rat Brain

**DOI:** 10.1101/2020.04.29.069542

**Authors:** Caitlin F. Fowler, Dan Madularu, Masoumeh Dehghani, Gabriel A. Devenyi, Jamie Near

**Affiliations:** Department of Biological and Biomedical Engineering, McGill University, Montreal, Canada; Centre d’Imagerie Cérébrale, Douglas Hospital Research Centre, Verdun, Canada; Center for Translational NeuroImaging, Northeastern University, Boston, MA, USA; Department of Psychiatry, McGill University, Montreal, Canada

**Author notes:** Corresponding author at: CIC Pavillion, office GH-2113, Douglas Hospital Research Centre, 6875, Boulevard LaSalle, Verdun, Canada, H4H 1R3.

**Keywords:** Proton Magnetic Resonance Spectroscopy, In Vivo, Aging, Biomarker, Macromolecule, Bioenergetics

## Abstract

Normal aging is associated with numerous biological changes including altered brain metabolism and tissue chemistry. *In vivo* characterization of the neurochemical profile during aging is possible using magnetic resonance spectroscopy, a powerful non-invasive technique capable of quantifying brain metabolites involved in physiological processes that become impaired with age. A prominent macromolecular signal underlies those of brain metabolites and is particularly visible at high fields; parameterization of this signal into components improves quantification and expands the number of biomarkers comprising the neurochemical profile. The present study reports, for the first time, the simultaneous absolute quantification of brain metabolites and individual macromolecules in aging male and female Fischer 344 rats, measured longitudinally using proton magnetic resonance spectroscopy at 7T. We identified age- and sex-related changes in neurochemistry, with prominent differences in metabolites implicated in anaerobic energy metabolism, antioxidant capacity, and neuroprotection, as well as numerous macromolecule changes. These findings contribute to our understanding of the neurobiological processes associated with healthy aging, critical for the proper identification and management of pathological aging trajectories.

**HIGHLIGHTS:**

→ Magnetic resonance spectroscopy reveals altered chemistry in the aging rat brain
→ Age- and sex-dependent differences in metabolites and macromolecules are present
→ Metabolites and macromolecules are markers of processes involved in healthy aging
→ This study clarifies normative progression of brain chemistry and metabolism *in vivo*
→ Improved understanding will inform future studies in pathological aging

## 1. INTRODUCTION

Aging is associated with impairments in numerous physiological pathways including antioxidant production, inflammatory response, and cerebral glucose metabolism, with further exacerbation seen in neurodegenerative disorders (Yin et al. 2016; Ravera et al. 2019; Miccheli et al. 2003). The medial temporal lobe, particularly the hippocampus, has a well-documented role in learning and memory consolidation, and is particularly vulnerable to the aging process and neurodegenerative disease (Bettio, Rajendran, and Gil-Mohapel 2017; Schuff et al. 1999; Van Hoesen, Hyman, and Damasio 1991). Improving our understanding of the neurobiological processes associated with healthy aging within the hippocampus may aid diagnosis, monitoring, and treatment of age-related neurological diseases such as dementia.

Proton magnetic resonance spectroscopy (^1^H-MRS) is a non-invasive technique that provides unique insight into brain metabolism *in vivo*, permitting longitudinal examination of the neurochemical profile. These neurochemicals can serve as biomarkers indicative of specific cellular and molecular mechanisms, in the context of both health and pathology (Ross and Sachdev 2004). For example, N-acetylaspartate (NAA) and myo-Inositol (Ins), are thought to reflect altered neuronal viability and gliosis, respectively, and have been reported to change over the course of normal aging (Haga et al. 2009; Cleeland et al. 2019; Harris et al. 2014), as well as in Alzheimer’s disease (Nilsen et al. 2014; Małgorzata Marjańska et al. 2019). MRS has been used to characterize altered tissue chemistry due to normal brain development and aging, disease, and therapeutic intervention in mice (Duarte, Do, and Gruetter 2014; M. Marjańska et al. 2014; Choi et al. 2014), rats (Paban, Fauvelle, and Alescio-Lautier 2010; Harris et al. 2014; Zhang et al. 2009), and humans (Murray et al. 2014; Emir et al. 2011), reflecting changes in underlying physiological processes and supporting its emergence as an important translational tool in neuroscience research

Quantification of at least 18 neurochemicals in the rodent brain is feasible with *in vivo* ^1^H MRS at 7T and above, providing a wide array of potential biomarkers of specific cellular and molecular changes (Pfeuffer et al. 1999; Mlynárik et al. 2006; Duarte, Do, and Gruetter 2014; Harris et al. 2014). Many of these molecular changes are reflective of changes in energy metabolism, inflammation, and antioxidant capacity, processes significantly altered in aging and whose metabolic by-products comprise the Nuclear Magnetic Resonance (NMR)-observable neurochemical profile (McKenna et al. 2012; Febo and Foster 2016). In addition to brain metabolites, broad macromolecule (MM) resonances are also detected with ^1^H MRS, and have been shown to change with age, brain region, and pathological conditions (Seeger et al. 2003; Behar et al. 1994; Hofmann et al. 2001). The MM peaks have previously been assigned to cytosolic proteins and mobile lipids (Behar and Ogino 1993), and contribute strongly to the overall signal (Považan et al. 2018; Cudalbu, Mlynárik, and Gruetter 2012).

Historically, the vast majority of MRS studies focus exclusively on quantification of metabolite levels, while only a few previous publications have sought to selectively measure the MM contribution. Generally, this has been achieved via inversion recovery (IR) experiments to suppress the metabolite signals. This metabolite-suppressed spectrum is then used as a basis to quantify the overall MM contribution in a standard MRS analysis. However, this approach does not allow for quantification of the individual MM components (Cudalbu, Mlynárik, and Gruetter 2012; Craveiro et al. 2014).

Alternatively, parameterization of the IR-derived MM signal into components has emerged as a viable option for determining the concentration of individual MMs; this process has been successfully applied to MRS and MRSI data obtained in humans from 1.5T and 7T (Seeger et al. 2003; Považan et al. 2018; Otazo et al. 2006), and in rats at 9.4T and 17.2T (Lee and Kim 2017; Lopez-Kolkovsky, Mériaux, and Boumezbeur 2016). Age-related change in MM concentration has been investigated in humans, but either the MM signal was quantified as a single entity (Małgorzata Marjańska et al. 2018), or individual MMs were quantified, but not longitudinally (Hofmann et al. 2001). To our knowledge, no preclinical studies have characterized longitudinal changes in individual MMs with age. Thus, the goal of this study was to assess longitudinal changes in both metabolites and individual macromolecules in the hippocampus of Fischer 344 rats to provide new insight into cellular mechanisms and biomarkers associated with healthy aging.

## 2. MATERIALS AND METHODS

### 2.1 Animals

Homozygous Fischer 344/NHsd wildtype (WT) male and female rats (henceforth referred to as Fischer) were obtained from Envigo Laboratories (Madison, WI, United States; order code: 010) and bred within the Animal Care Facility at the Douglas Hospital Research Centre to avoid confounds from the stress of transportation. Rats were weaned on postnatal day 21 and housed in pairs on a 12 hour light-dark cycle with *ad libitum* access to food and water. All animal procedures and experiments were performed in accordance with the guidelines of the McGill University Animal Care Committee and the Douglas Hospital Research Centre Animal Care Committee.

Neurochemical profiles were measured longitudinally, with scans at 4-, 10-, 16-, and 20-months of age, covering the majority of the adult rat lifespan. A total of 30 rats were included in the study. 12 of 30 rats were scanned at only the 4- and 10-month timepoints due to being part of a separate treatment study thereafter. Additionally, 2 females died prior to the 16-month time point and 5 males died prior to the 20-month time point, all of natural causes, and one scan at 10-months was discarded due to low SNR. As such, the final number of scans for the four time points were 30 (15 males, 15 females), 29 (14 males, 15 females), 16 (7 males, 9 females), and 11 (3 males, 8 females), respectively (**see Supplementary Table 1**). The imbalance in the number of subjects scanned per time point was accounted for in the statistical analysis by applying a mixed effects linear model.

### 2.2 ^**1**^H-MRS data acquisition - metabolites

MRS data were acquired at the Douglas Centre d’Imagerie Cérébrale using a 7 Tesla Bruker Biospec 70/30 scanner (Bruker, Billerica, MA, United States) with an 86 mm volumetric birdcage coil for transmission and a four-channel surface array coil for signal reception (Bruker). The level of anesthesia (1- 4% isoflurane in oxygen gas) was adjusted to maintain a breathing rate between 50-70 breaths per minute throughout the procedure and warm air (37 °C) was blown into the bore of the scanner to maintain a constant body temperature (SA Instruments, Inc., monitoring system, Stony Brook, NY, United States).

Coronal and sagittal T_2_-weighted MR images were acquired using a Rapid Acquisition with Relaxation Enhancement (RARE) sequence (TR = 2049 ms, TE = 10.8 ms, RARE factor = 6, effective echo time = 32.4 ms, FOV = 32 x 22.5 mm, matrix size = 256 x 180, resolution = 125 μm non-isotropic, 1m26s70ms acquisition time) to guide placement of a region of interest for localized magnetic resonance spectroscopy in the dorsal hippocampus (∼31 μL volume, see **Figure 1**). Voxel positioning was based on anatomic landmarks. All spectra were acquired from the right hemisphere.

**Figure 1.**
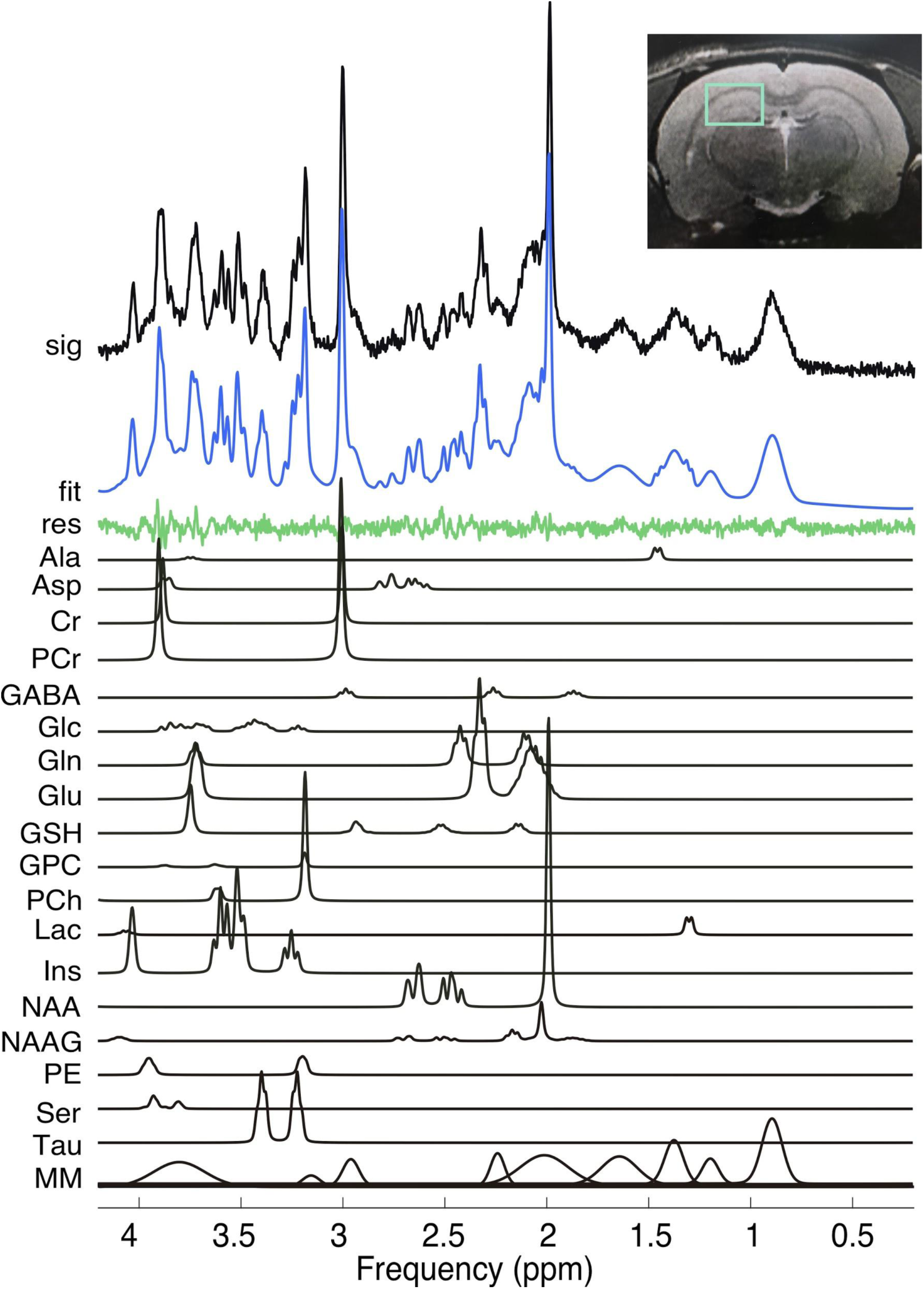
Localized 1H-MRS spectra from the hippocampus of a 16-month old female rat acquired using PRESS at 7T. The spectrum was processed using the FID-A toolkit which includes removal of bad averages, frequency drift correction, and zero-order phasing using the creatine peak. Metabolites included in the basis set are shown and assigned as follows: Ala, alanine; Asp, aspartate; Cr, creatine; PCr, phosphocreatine; GABA, gamma-aminobutyric acid; Glc, glucose; Gln, glutamine; Glu, glutamate; GSH, glutathione; GPC, glycerophosphocholine; PCh, phosphocholine; Lac, lactate; Ins, myo-inositol; NAA, N-acetylaspartate; NAAG, N-acetylaspartyl glutamate; PE, phosphoethanolamine; Ser, serine; Tau, taurine; MM, macromolecule. **Top right:** a representative RARE image shows the typical placement of the volume of interest for spectroscopy

Automated localized shimming was performed using the FASTMAP method (Gruetter 1993) (ParaVision 5.1, Bruker). Specifically, 1st and 2nd order shims were first optimized on a 5×5×5 mm^3^ voxel, followed by 1st order-only shimming on a smaller, local 3.5×3.5×3.5 mm^3^ voxel (average water linewidth 9.92 ± 1.03 Hz), both surrounding the region of interest. *In vivo* ^1^H-MRS scans were then acquired from a 2.5×3.5×3.5 mm^3^ voxel using the PRESS sequence (Point Resolved Spectroscopy) with a total acquisition time of 13m0s0ms (TR = 3000 ms, TE = 11.12 ms, 2048 acquisition data points, spectral width = 4006 Hz) in combination with outer volume suppression. 256 averages were acquired with VAPOR water suppression (Tkač et al. 1999) and 8 averages were acquired without water suppression for frequency, phase shift, and eddy current correction, as well as absolute concentration calculations, described in the **Supplementary Methods**.

### 2.3 ^**1**^H-MRS data acquisition - macromolecules

“Metabolite-nulled” spectra were acquired in eight Fischer rats at 10-months of age using PRESS localization (TR = 3000 ms, TE = 11.12 ms, 512 averages, 2048 acquisition data points, spectral width = 4006 Hz) preceded by an IR module consisting of a 4th order hyperbolic secant adiabatic full passage (AFP) inversion pulse (pulse duration =1.0 ms, bandwidth = 6982.0 Hz, inversion time = 800 ms), for a total acquisition time of 25m48s0ms. The eight macromolecule spectra were aligned, summed, and scaled in FID-A to generate an average metabolite-nulled spectrum for parameterization. For details on spectral processing in FID-A, see **Supplementary Methods**. Due to variability in longitudinal relaxation times of metabolites, a single inversion time is unlikely to produce a macromolecule spectrum wherein all metabolites are completely suppressed(Cudalbu, Mlynárik, and Gruetter 2012). To identify the residual metabolite peaks in our MM spectrum, two additional MM scans were obtained with the same inversion time (TI=800 ms) and scan parameters as above, but with a long TE (40 ms). Minor residual peaks of NAA (2.00 ppm, inverted; 2.69 ppm), tCr (3.93 ppm), Glu (2.36 ppm), Gln (2.53 ppm), and Taurine (3.41 ppm, inverted) were visible and accounted for during the parameterization process.

### 2.4 Parameterization of the macromolecule spectrum

Parametrization of the MM spectrum was performed in jMRUI using the AMARES fitting tool(Vanhamme, van den Boogaart A, and Van Huffel S 1997; Stefan et al. 2009; Naressi et al. 2001). The water peak was manually removed from the averaged MM spectrum using HLSVD (Hankel-Lanczos singular value decomposition(Pijnappel et al. 1992)), in jMRUI and first order phasing was performed. Then, prior knowledge of the chemical shifts of ten MM resonances of interest was used as the input in the initial phase of modeling using Gaussian functions, wherein the subscript denotes the frequency at which the resonance appears: MM_0.89_, MM_1.20_, MM_1.39_, MM_1.66_, MM_2.02_, MM_2.26_, MM_2.97_, MM_3.18_, MM_3.84_, and MM_4.27(Považan et al. 2018; Otazo et al. 2006; Lee and Kim 2017)_. The MM peak at 4.27 ppm was parameterized but later omitted from basis set simulations due to proximity to the water peak.

As suggested by Craveiro, et al., residual metabolite signals within the MM spectrum can be better accounted for by including advanced prior knowledge of the residual peaks in the fitting process(Craveiro, Cudalbu, and Gruetter 2012; Craveiro et al. 2014). Constraints were therefore manually set for the frequency, phase, and linewidth of residual metabolite peaks (NAA, tCr, Glu, Gln, and Tau), and modelled using Lorentzian functions. Soft constraints on the overall and relative phase, frequency, and linewidth of all peaks were modified iteratively to achieve minimal fitting residuals.

### 2.5 Generation of metabolite and macromolecule basis spectra

The FID-A Simulation toolbox (github.com/CIC-methods/FID-A, version 1.0, (Simpson et al. 2017) in MATLAB (R2012a, The MathWorks, Inc., Natick, Massachusetts, United States) was used to simulate all metabolite and macromolecule basis functions for LCModel analysis. Simulations took into account the exact PRESS refocusing pulse waveforms that were used experimentally (Mao, Mareci, and Andrew 1988), as well as the actual TE and TR, and metabolite spin systems were based on previously published chemical shifts and J-coupling constants (Govindaraju, Young, and Maudsley 2000). Metabolites were simulated with Lorentzian lineshapes and linewidths of 2 Hz, while macromolecules were simulated using Gaussian lineshapes, while linewidths and frequency were based on jMRUI AMARES output (see **Supplementary Table 2** for MM basis set simulation parameters). The neurochemical basis set consisted of 18 simulated metabolite resonances and 9 macromolecule basis functions: alanine (Ala), aspartate (Asp), creatine (Cr), γ-aminobutyrate (GABA), glucose (Glc), glutamine (Gln), glutamate (Glu), glycerophosphocholine (GPC), glutathione (GSH), lactate (Lac), myo-Inositol (Ins), N-acetylaspartate (NAA), N-acetylaspartylglutamate (NAAG), phosphocholine (PCh), phosphocreatine (PCr), phosphoethanolamine (PE), serine (Ser), taurine (Tau), MM_0.89_, MM_1.20_, MM_1.39_, MM_1.66_, MM_2.02_, MM_2.26_, MM_2.97_, MM_3.18_, MM_3.84_.

### 2.6 Data processing and quantification

All raw proton MRS spectra were processed using the FID-A Processing toolbox, which enables the removal of motion corrupted scans and correction of frequency and phase drift errors. Spectra were analyzed using the linear combination analysis method LCModel (version 6.3, Stephen Provencher Inc, Oakville, Ontario, Canada) using basis functions simulated in FID-A. Soft constraints on concentration ratios were specified for individual MM resonances, based on AMARES amplitudes (see **Supplementary Table 2**). Fitting was performed over the spectral range of 0.2 to 4.2 ppm. The unsuppressed water signal measured from the same volume of interest was used as an internal reference for absolute quantification.

The detailed output from LCModel was used to construct a matrix of correlation coefficients between the metabolite concentrations across all subjects. If the average correlation between a pair was less than -0.3, we chose to also report those metabolites as a sum (S. W. Provencher 1993; S. Provencher 2019). Therefore, we included the following sums in our neurochemical profile: GPC+PCh (total Choline, tCho), Cr+PCr (total creatine, tCr), and Tau+Glc. We also report summed NAA+NAAG (tNAA) and Glu+Gln (Glx) and the ratios of PCr to Cr, Glu to Gln, Asp to Glu, and NAA to Ins.

### 2.7 Application of correction factors for absolute quantification

For absolute quantification of metabolites and macromolecules, we applied a correction factor to the raw water referenced data provided by LCModel. This correction factor was determined using an equation modified from Dhamala et al. (Dhamala et al. 2019), and accounted for T1 and T2 relaxation constants of water and measured neurochemicals, as well as a correction factor to account for the fact that our voxel contained primarily grey matter (with an estimated NMR-visible water concentration of 43300 mM, (Ernst, Kreis, and Ross 1993). See **Supplementary Equation 2** and **Supplementary Table 3** for the final correction factors applied to the LCModel concentration outputs for all neurochemicals. All neurochemical concentrations are reported in mmol/L.

### 2.8 Exclusion criteria

The Cramer-Rao lower bound (CRLB) provided by LCModel was used as a measure of reliability of neurochemical quantification on a per-metabolite basis. CRLBs (or %SD) are estimates of the uncertainty of the concentrations calculated by LCModel and are expressed in percent of the estimated concentrations. We chose to use a strict cut-off of 30% CRLB averaged across all scans, which resulted in the removal of GABA, Serine, and MM_3.18_ from our analysis. See **Supplementary Table 4** for details on metabolite CRLB across the four time points. These exclusion criteria are based on recommendations by Kreis et al., wherein only entire metabolites are to be excluded, as opposed to excluding specific samples with low % SD, in order to avoid biasing the mean estimated concentrations of cohort data (Kreis 2016).

### 2.9 Statistical analysis

Metabolites were analyzed using linear mixed effects modeling in R (version 3.6.3, tidyverse_1.3.0, lme4_1.1-21, lmerTest_3.1-0) with the fixed effect variables defined by age and sex, and rat ID as a random effect. Our study was underpowered to assess the presence of an age by sex interaction so a test for a main effect of sex was performed by collapsing across the 4 time points. The fixed effect of water linewidth (water.lw), extracted from the water unsuppressed data using the FID-A toolkit in MATLAB, was included to control for the effect of linewidth on metabolite concentration estimates (Bartha 2007). A weighting factor of the inverse absolute standard deviation (or CRLB) for each metabolite (metabolite.sdab) was included to account for differences in fitting reliability between samples, and allowed us to include all observations with CRLB <999. All continuous variables were z-scored so that coefficients (betas, or effect sizes) for each fixed effect would be standardized. As such, the standard betas indicate how many standard deviations metabolite concentration has changed per standard deviation increase in the predictor variable (age or sex). The full linear model is shown below.

**Equation 1:** lmer(scale(*metabolite*))∼scale(*age*)+s*ex*+scale(*water*.*lw*)+(*1*|*subject)*, weights = 1/scale(*metabolite*.*sdab*)

Due to the number of comparisons, we used the False Discovery Rate method (Benjamini and Hochberg 1995) to control the family-wise type I error rate at 5% for the main effects of age and sex. Both p- and q- (or FDR-corrected) values are presented in **Table 1**. A fixed effect was considered significant when q < 0.05.

**Table 1.** Change in neurochemical concentrations with age and sex in Fischer rats. Std.Beta. represents the standardized beta or coefficient value for the variable of sex or age in the linear model used for analysis. q.value is the FDR-corrected p-value obtained for each fixed effect. P and q <0.05 are denoted in bold. Abbreviations: Ala, alanine; Asp, aspartate; Cr, creatine; PCr, phosphocreatine; GABA, gamma-aminobutyric acid; Glc, glucose; Gln, glutamine; Glu, glutamate; GSH, glutathione; GPC, glycerophosphocholine; PCh, phosphocholine; Lac, lactate; Ins, myo-inositol; NAA, N-acetylaspartate; NAAG, N-acetylaspartyl glutamate; PE, phosphoethanolamine; Ser, serine; Tau, taurine; tCho, total choline (GPC+PCh); tCr, total creatine (Cr+PCr); Glx (Glu+Gln); MM, macromolecule.

|  | Main effect of age |  |  |  | Main effect of sex |  |  |  |
| --- | --- | --- | --- | --- | --- | --- | --- | --- |
|  | Std. Beta | p.value | q.value | Power (%) | Std. Beta | p.value | q.value | Power (%) |
| Ala | -0.092 | 0.387 | 0.511 | 15.90 | -0.289 | 0.228 | 0.329 | 21.70 |
| Asp | -0.222 | **0.040** | 0.080 | 53.50 | 0.108 | 0.650 | 0.733 | 5.60 |
| Cr | 0.147 | 0.152 | 0.236 | 29.30 | -0.406 | 0.106 | 0.177 | 32.20 |
| PCr | -0.231 | **0.022** | 0.062 | 60.10 | 0.817 | **0.001** | **0.005** | 92.30 |
| Glc | 0.324 | **0.001** | **0.005** | 92.10 | -1.129 | **<0.0001** | **0.001** | 99.40 |
| Gln | 0.072 | 0.497 | 0.616 | 10.00 | -0.383 | 0.144 | 0.228 | 32.30 |
| Glu | -0.003 | 0.978 | 0.978 | 5.40 | 0.379 | 0.084 | 0.149 | 37.00 |
| GSH | -0.335 | **0.001** | **0.006** | 91.10 | 0.331 | 0.174 | 0.263 | 24.60 |
| Lac | 0.355 | **0.001** | **0.005** | 92.70 | -0.033 | 0.886 | 0.931 | 4.10 |
| Ins | 0.363 | **0.001** | **0.005** | 91.90 | -0.022 | 0.930 | 0.961 | 3.70 |
| NAA | -0.005 | 0.959 | 0.975 | 5.90 | 0.444 | 0.067 | 0.121 | 40.50 |
| NAAG | 0.329 | **0.001** | **0.006** | 89.50 | -0.479 | **0.035** | 0.077 | 52.60 |
| PE | -0.214 | **0.038** | 0.080 | 54.60 | -0.500 | **0.034** | 0.077 | 55.60 |
| Tau | -0.244 | **0.023** | 0.062 | 63.00 | -0.177 | 0.489 | 0.616 | 10.70 |
| tCho | 0.089 | 0.340 | 0.458 | 15.40 | 0.473 | **0.028** | 0.067 | 53.40 |
| tCr | -0.176 | 0.094 | 0.162 | 38.20 | 0.652 | **0.006** | **0.020** | 73.90 |
| tNAA | 0.064 | 0.537 | 0.653 | 9.70 | 0.358 | 0.124 | 0.202 | 29.60 |
| Glx | 0.022 | 0.823 | 0.891 | 6.80 | 0.172 | 0.474 | 0.613 | 9.30 |
| Tau+Glc | 0.120 | 0.210 | 0.309 | 23.80 | -1.082 | **<0.0001** | **0.001** | 99.30 |
| Glu/Gln | -0.060 | 0.561 | 0.669 | 8.50 | 0.551 | **0.025** | 0.064 | 58.40 |
| PCr/Cr | -0.216 | **0.021** | 0.062 | 59.30 | 0.659 | **0.004** | **0.015** | 77.70 |
| NAA/Ins | -0.254 | **0.006** | **0.020** | 79.00 | 0.480 | **0.040** | 0.080 | 52.60 |
| Asp/Glu | -0.243 | **0.026** | 0.064 | 60.20 | -0.055 | 0.820 | 0.891 | 5.20 |
| MM<sub>0.89</sub> | 0.331 | **0.001** | **0.005** | 92.00 | 0.097 | 0.645 | 0.733 | 5.70 |
| MM<sub>1.20</sub> | 0.380 | **<0.0001** | **0.0003** | 99.80 | -0.654 | **0.003** | **0.013** | 87.70 |
| MM<sub>1.39</sub> | 0.113 | 0.249 | 0.351 | 21.90 | 0.592 | **0.012** | **0.038** | 70.00 |
| MM<sub>1.66</sub> | 0.357 | **0.0005** | **0.005** | 92.20 | 0.050 | 0.834 | 0.891 | 4.80 |
| MM<sub>2.02</sub> | 0.423 | **<0.0001** | **<0.0001** | 99.90 | 0.593 | **0.001** | **0.006** | 88.20 |
| MM<sub>2.26</sub> | 0.177 | 0.060 | 0.114 | 47.30 | 0.459 | 0.060 | 0.114 | 47.50 |
| MM<sub>2.97</sub> | -0.051 | 0.600 | 0.702 | 8.20 | 0.792 | **0.003** | **0.013** | 86.60 |
| MM<sub>3.84</sub> | 0.290 | **0.003** | **0.011** | 86.90 | -0.226 | 0.313 | 0.431 | 15.00 |

Finally, observed power analysis was performed for each metabolite using the CRAN package, SIMR (R version 3.63, simr_1.0.5, (Green and MacLeod 2016) to ensure our results from a relatively small sample size were generalizable to a larger population. See **Supplementary Methods** for details.

## 3. RESULTS

### 3.1 Quantification using an expanded neurochemical profile

A representative NMR spectrum from a female rat brain at 16-months shows the excellent metabolite spectral quality consistently obtained in this study (**Figure 1**). Spectral linewidth across timepoints, measured from the unsuppressed water signal, varied slightly (t1: 9.73 ± 0.94 Hz, t2: 10.38 ± 1.26 Hz), t3: 9.64 ± 0.66 Hz, t4: 9.60 ± 0.56 Hz). All linewidths were well under the 0.1 ppm (30.3 Hz at 7T) full-width half-max (FWHM) considered essential for in vivo 1H MRS spectra (Forster 2012). More detail on spectral linewidth at each timepoint, as well as SNR, can be found in **Supplementary Table 5**.

**Figure 1**. also shows the 18 metabolite and nine macromolecule basis functions used by LCModel to fit each spectrum. With the exclusion of high-CRLB metabolites GABA, Ser, and MM_3.18_, the following compounds comprised the neurochemical profile used to evaluate changes with age and differences between sexes: Ala, Asp, Cr, PCr, Glc, Gln, Glu, GSH, Ins, Lac, NAA, NAAG, PE, Tau, tCho (GPC+PCh), tCr (Cr+PCr), Glx (Glu+Gln), tNAA (NAA+NAAG), Tau+Glc, Glu/Gln, PCr/Cr, NAA/Ins, Asp/Glu, MM_0.89_, MM_1.20_, MM_1.39_, MM_1.66_, MM_2.02_, MM_2.26_, MM_2.97_, MM_3.18_, and MM_3.84_. All 31 neurochemicals can be said to have been quantified with high reliability: 20/31 and 30/31 neurochemicals displayed average CRLBs of < 10% and < 20%, respectively, with only Alanine, at 21.5%, displaying a CRLB greater than 20%. See **Supplementary Table 4** for details on metabolite CRLB.

The average metabolite-nulled MM spectrum and its parameterization into nine components in AMARES is shown in **Figure 2**. The amplitudes and line widths of the MM peaks determined in this step were used to create the MM basis functions for LCModel analysis. **Figure 2**. also shows a representative spectrum without and with the individual MMs in the basis set. The improved fit quality upon incorporation of MM resonances is reflected by the flatness of the resulting spline baseline, which represents the tail of the residual water peak.

**Figure 2.**
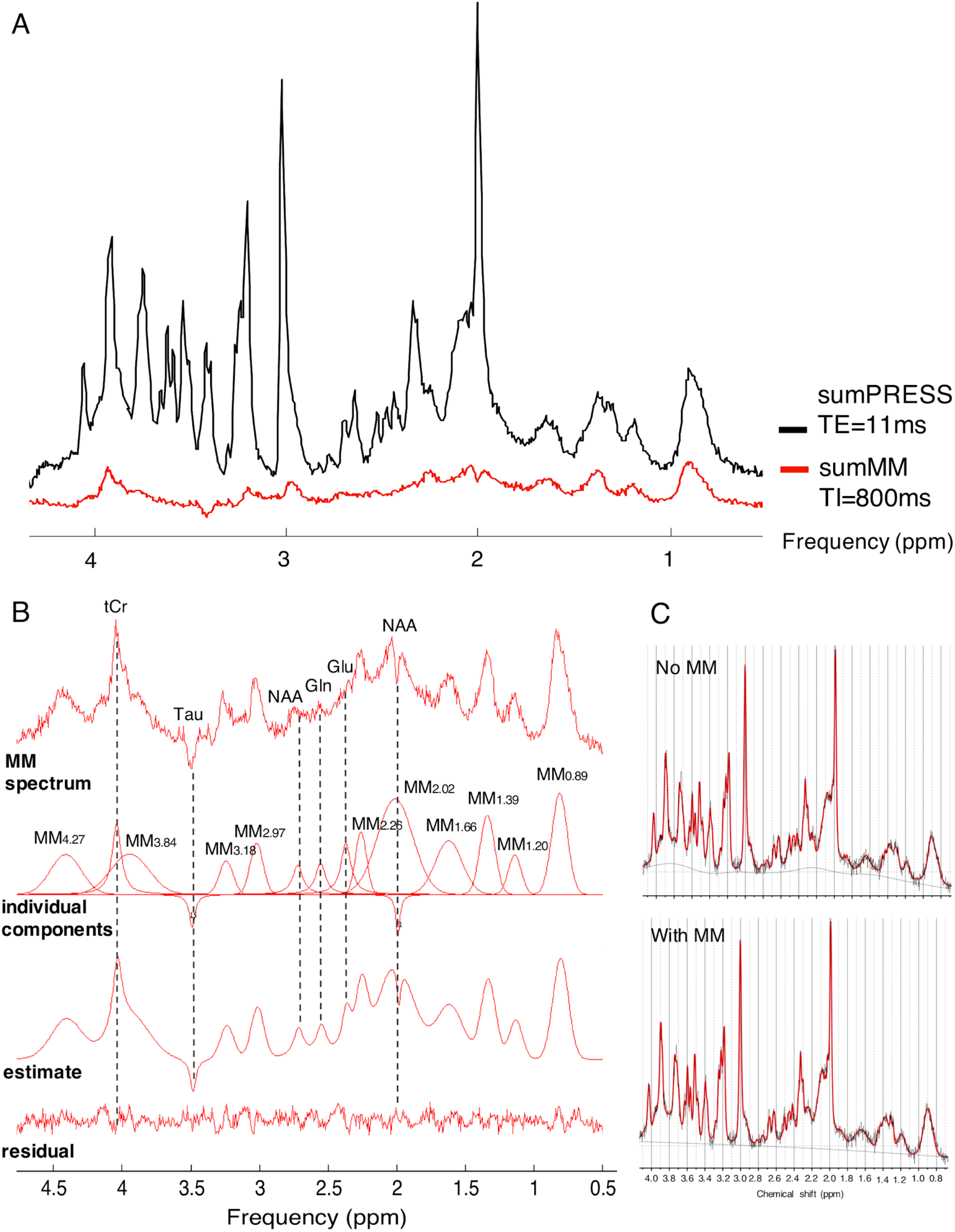
Parameterization of the metabolite-suppressed spectrum. (A) Average localized ^1^H-NMR spectra (n=8, PRESS sequence) and corresponding metabolite-suppressed spectra (n=8, Inversion Recovery PRESS sequence) used for parameterization. (B) Output of AMARES quantification showing the average metabolite-nulled spectra with all individual components used for fitting (including residual metabolite peaks), the overall fitting estimate, and the fit residual). (C) A representative spectrum analyzed in LCModel with the default macromolecule basis functions (top), versus the same spectrum analyzed in LCModel using the basis functions simulated using prior knowledge from the parameterized MM spectral data (bottom). The improved fit quality upon incorporation of MM resonances in the basis set is reflected by the flatness of the resulting spline baseline in the bottom frame.

Regarding neurochemical concentration, all metabolite concentrations were well within ranges previously reported in the rat brain by numerous authors whose quantification methods incorporated MMs (Harris et al. 2014; Lopez-Kolkovsky, Mériaux, and Boumezbeur 2016; Pfeuffer et al. 1999). As individual MMs have not yet been quantified in rodent brain, we compared our macromolecule concentrations to those cited in a study by Snoussi et al, wherein MMs in healthy adults were quantified in the range of 5-20 mmol/kg, which is in good agreement with our findings (Snoussi et al. 2015).

### 3.2 Metabolite and macromolecule concentrations are altered with age

The levels of 11 of 31 neurochemicals changed significantly with age in Fischer rats (**Table 1**), using q<0.05. Change in concentration of metabolites, metabolite ratios, and macromolecules with age are shown in **Figures 3 and 4**. We observed significant concentration reductions with age for GSH (β = - 0.335) only. Increasing concentrations with age were seen for Glc (β=0.324), Lac (β = 0.355), Ins (β = 0.363), NAAG (β = 0.329), MM_0.89_ (β = 0.331), MM_1.20_ (β = 0.380), MM_1.66_ (β = 0.357), MM_2.02_ (β = 0.423), and MM_3.84_ (β = 0.290). Of the four metabolite ratios analyzed, only NAA/Ins was significantly altered with age (β = -0.254). Absolute concentrations of each metabolite at each timepoint, along with the q-value and standardized beta for the effect of age, are shown in **Supplementary Table 6**.

**Figure 3.**
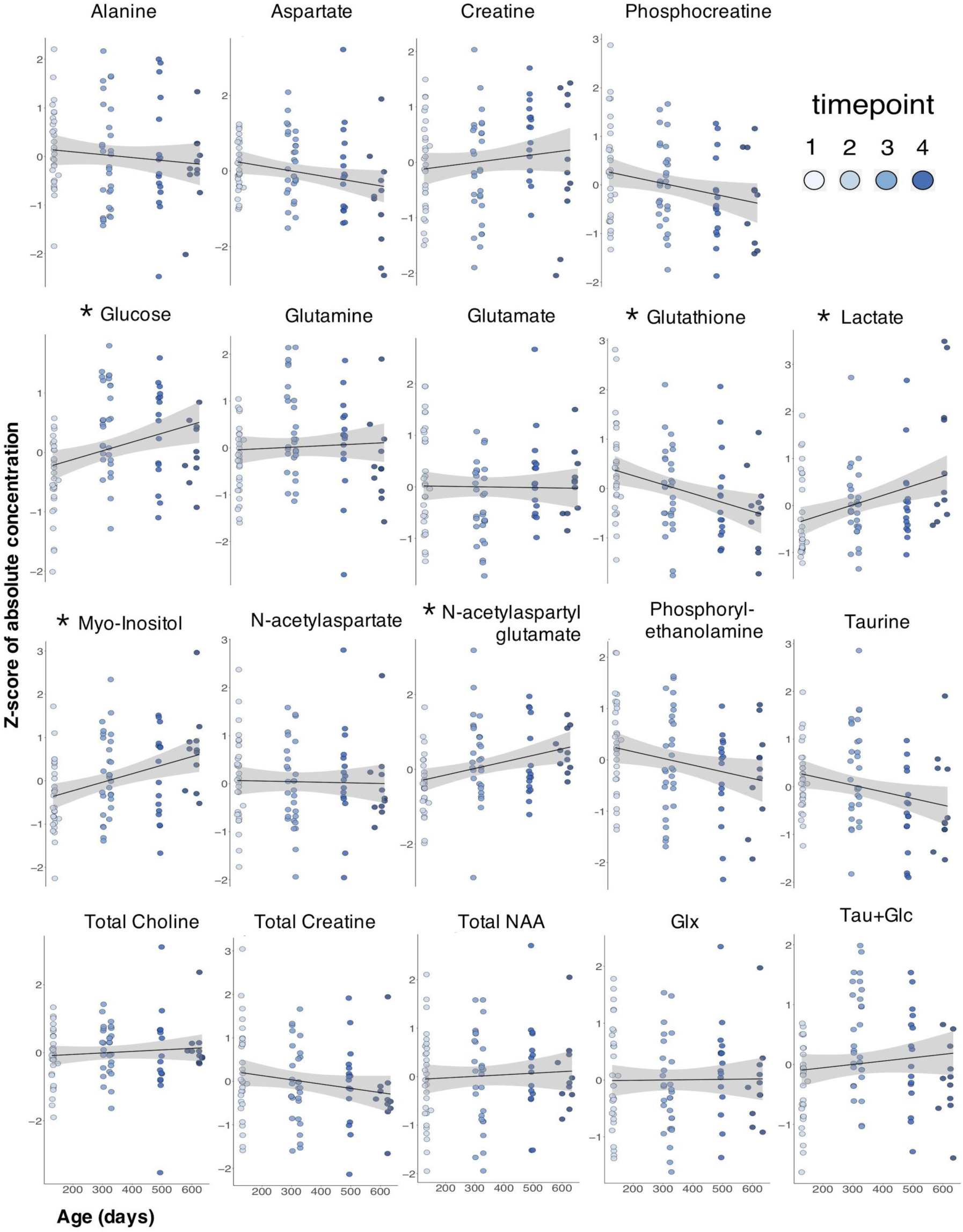
Age-dependent change in metabolite concentrations in the hippocampus of Fischer rats. ^1^H-NMR spectra were acquired longitudinally in rats aged 4-, 10-, 16-, and 20-months old, and an effect of age was determined using linear mixed effects modelling with FDR correction. * represents significant effects of age at q < 0.05.

**Figure 4.**
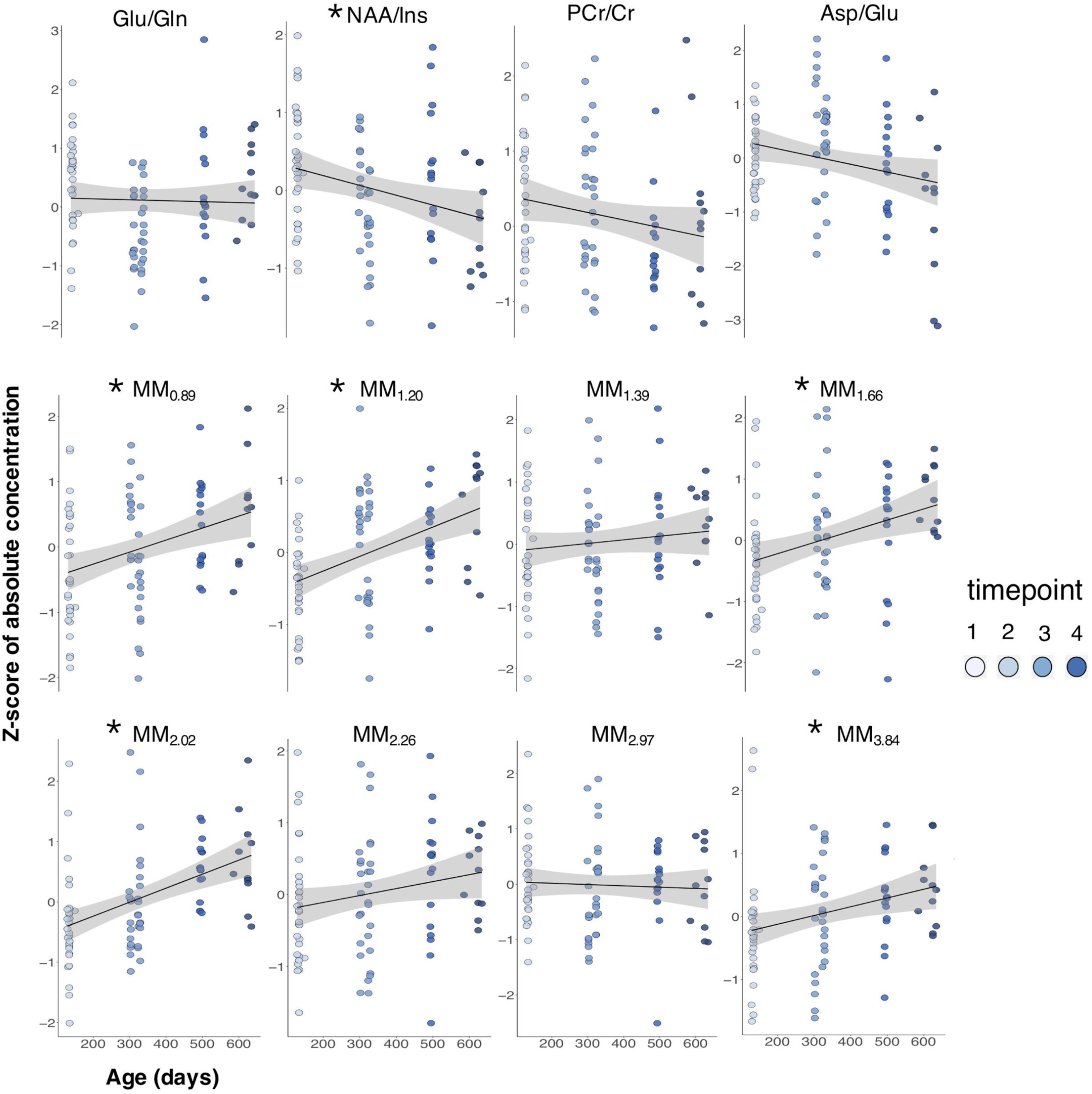
Age-dependent change in brain metabolite ratios and macromolecules in Fischer rats. ^1^H-NMR spectra were acquired longitudinally in rats aged 4-, 10-, 16-, and 20-months old, and an effect of age was determined using linear mixed effects modelling with FDR correction. * represents significant effects of age at q < 0.05.

### 3.3 Male and female Fischer rats exhibit differences in brain tissue chemistry

9 of 31 neurochemicals showed significant differences between male and female Fischer rats when examining data collapsed across all timepoints, using q<0.05 (**Table 1**). Differences in concentration of metabolites, metabolite ratios, and macromolecules due to sex are shown in **Figures 5 and 6**, respectively. The concentration of PCr was significantly increased in females relative to males (β=0.817), as was tCr (β=0.652), the ratio of PCr/Cr (β=0.659), MM_1.39_ (β=0.592), MM_2.02_ (β=0.593) and MM_2.97_ (β=0.792). Glc (β=-1.129), Tau+Glc (β=-1.082), and MM_1.20_ (β=-0.654), were decreased in females relative to males, collapsed across all timepoints. Absolute concentrations of each metabolite collapsed across timepoints and split by sex, along with the q-value and standardized beta for the main effect of sex, are shown in **Supplementary Table 7**.

**Figure 5.**
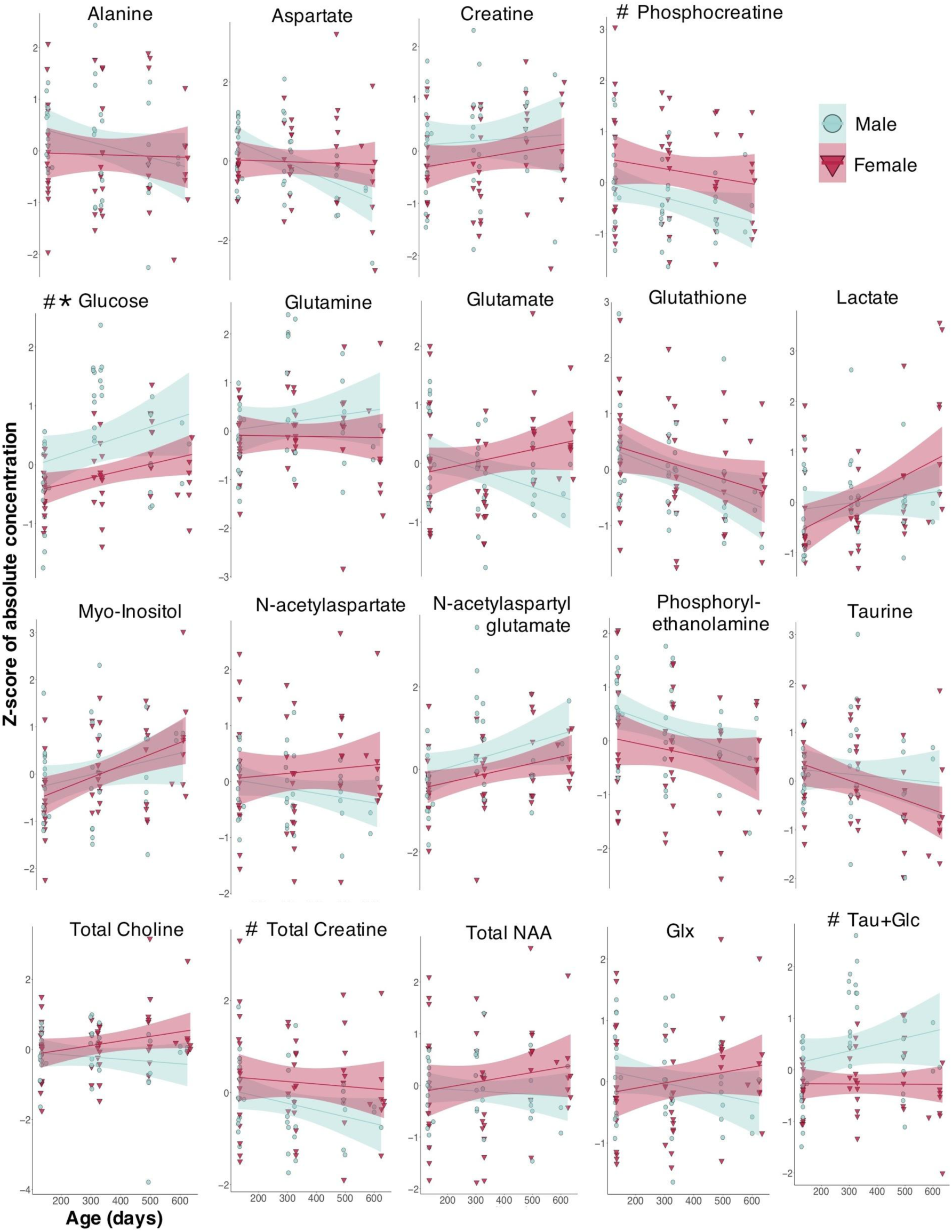
Differences in hippocampal metabolite concentrations exist between male (teal points) and female (red points) Fischer rats. ^1^H NMR spectra were acquired ongitudinally in rats aged 4-, 10-, 16-, and 20-months old, and a main effect of sex was determined using linear mixed effects modelling with FDR correction using data collapsed across all four timepoints. * and # represents significant effects of age and sex, respectively, at q < 0.05.

**Figure 6.**
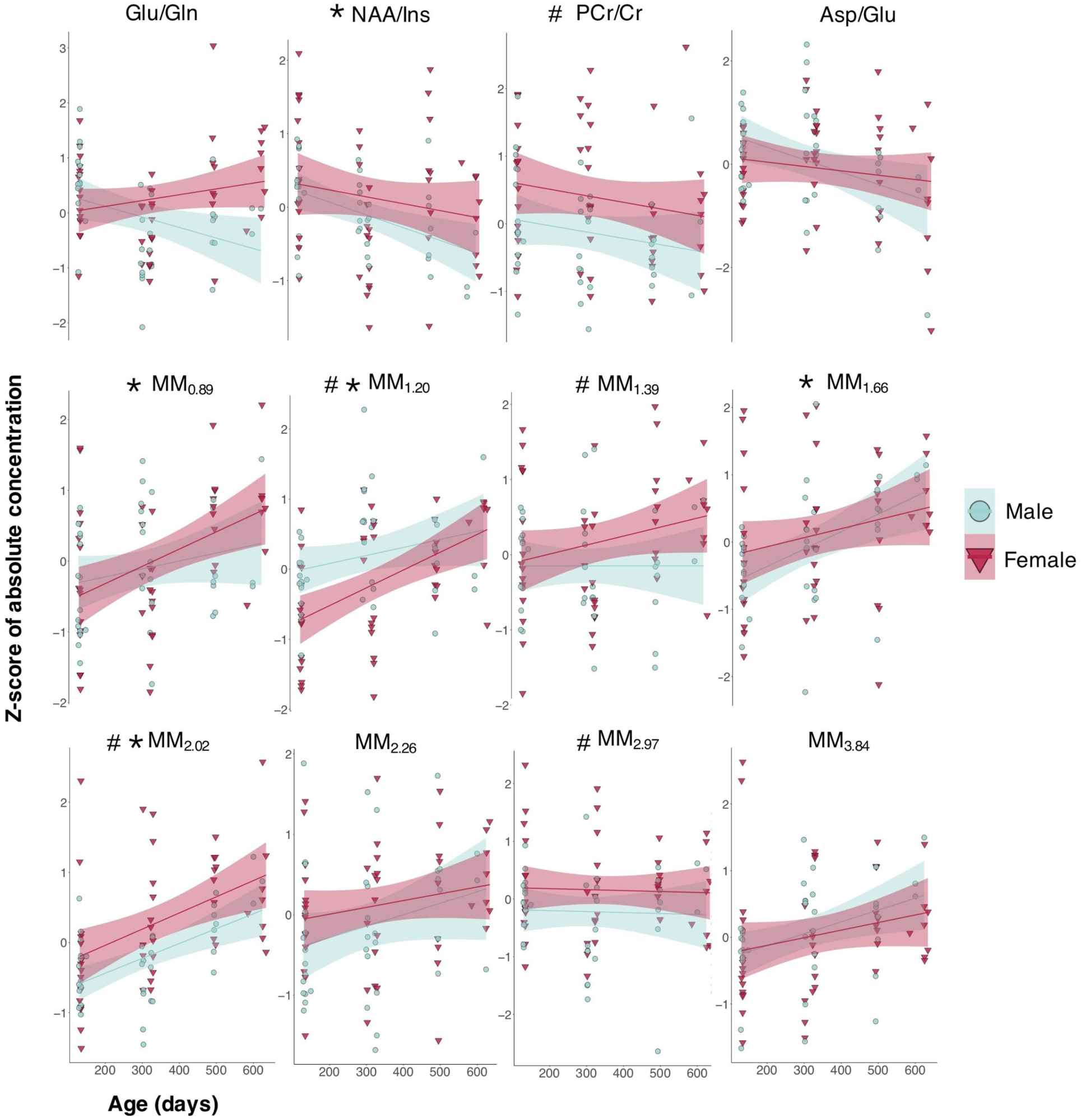
Differences in brain metabolite ratios and macromolecule concentrations exist between male (teal points) and female (red points) F344 rats. ^1^H NMR spectra were acquired longitudinally in rats aged 4-, 10-, 16-, and 20- months old, and a main effect of sex was determined using linear mixed effects modelling with FDR correction using data collapsed across all four timepoints. * and # represents significant effects of age and sex, respectively, at q < 0.05.

### 3.4 Age- and sex-dependent changes in neurochemistry over three timepoints versus four timepoints

Finally, to ensure our results were not being driven by the last time point wherein we have the fewest subjects (and therefore, the least power), we also analyzed all data from time points 1 through 3. Of the eleven neurochemicals with age-related changes over the four timepoints, four (Glc, GSH, MM2, MM5) remained significant at a q-value of < 0.05, five (Ins, NAAG, MM_0.89_, MM_1.66_, MM_3.84_) were significant at a p-value of < 0.05 and trending at a q-value of ≤ 0.1, one (NAA/Ins) had a p-value < 0.1 but a q-value > 0.1, and one (Lac) was no longer significant, with p-value and q-values of 0.105 and 0.181, respectively. Of the nine neurochemicals with significant differences (q < 0.05) between males and females, eight remained significant and the last was trending (p<0.1) when analyzed using only the 4-, 10-, and 16-month data. For more details see **Supplementary Tables 8 and 9**.

## 4. DISCUSSION

The present study reports, for the first time, the simultaneous absolute quantification of metabolites and individual macromolecules in the aging Fischer rat brain, measured longitudinally using high field ^1^H-MRS. The addition of modelled macromolecule resonances to the standard array of metabolites measured by ^1^H-MRS not only significantly improves metabolite quantification, but also expands the number of potential biomarkers available for detecting pathological changes in brain tissue metabolism (Hofmann et al. 2002). Using this expanded basis set of 31 neurochemicals, we identified age- and sex-related changes in tissue chemistry in the hippocampus, a brain region with a well-documented role in the cognitive decline that occurs with aging (Schuff et al. 1999; Bettio, Rajendran, and Gil-Mohapel 2017; Morrison, Stern, and Carstensen 2000).

### 4.1 Comparison with Previous Studies

Previous attempts at modelling MM components in proton MRS spectra have been made. In 2003, Seeger et al. first proposed parameterizing a metabolite-nulled spectrum for inclusion into the quantification basis set (Seeger et al. 2003); they adequately modelled the MM contribution in human brain spectra acquired at 1.5T using only four broad MM components, based on resonances identified by Behar in 1994 (Behar et al. 1994). As MM contributions become more resolved at higher field strengths, additional components and more strict fitting constraints are needed to prevent overestimation of overlapping metabolite concentrations (Cudalbu, Mlynárik, and Gruetter 2012; Pfeuffer et al. 1999; Považan et al. 2018).

In recently published human MRSI data at 7T by Povazan et al. employed direct spectral fitting of nine individual MMs as part of the basis set and revealed spectroscopic differences between grey matter (GM) and white matter (WM) in healthy adults (Považan et al. 2018). The MM profile of healthy adults has also been examined via single-voxel MRS at 3T and 7T; Snoussi et al. reported concentration values in the range of 5-20 mmol/kg for seven MM components fit using the sum of 19 Gaussian line shapes (Snoussi et al. 2015). Single-voxel animal studies have revealed that a more complex MM baseline can be resolved at higher field strengths, necessitating additional model components and constraints. Authors Lee and Kim proposed the direct parameterization of the MM baseline directly from short echo time spectra acquired at 9.4 T in rat brain using 25 components (Lee and Kim 2017), while Lopez-Kolkovsky et al used 32 individual resonances to parameterize metabolite-nulled rat brain spectra acquired at 17.2T (Lopez-Kolkovsky, Mériaux, and Boumezbeur 2016).

In the rat brain at 7T, nine to ten distinct macromolecular resonances are visible, similar to those identified in the human brain at 7T by Povazan and Snoussi (Považan et al. 2018; Snoussi et al. 2015). Although preclinical studies are attractive due to the expanded metabolic profile available at high fields and the large number of available disease models, little work has been done regarding preclinical quantification of both metabolites and MMs with age and between sexes.

### 4.2 Change in metabolite concentrations associated with healthy aging

Considering the established relationship between cognitive decline and decreased hippocampal structure and integrity - both in the context of aging and neurodegenerative disease (Bettio, Rajendran, and Gil-Mohapel 2017; Van Hoesen, Hyman, and Damasio 1991) - we chose to characterize the neurochemical profile of the aging hippocampus. Deficits in cognitive function with age are associated with impaired cerebral glucose metabolism and energy balance, with further exacerbation seen in neurodegenerative disorders (Yin et al. 2016; Ravera et al. 2019; Miccheli et al. 2003). Specifically, aging induces a shift from aerobic to anaerobic energy metabolism, due in part to decreased neuronal glucose uptake and mitochondrial energy-transducing capacity, and resulting in increased oxidant production, decreased tricarboxylic acid cycle (TCA) activity, and compromised electron transport and oxidative phosphorylation activity (Yin et al. 2016; Dong and Brewer 2019; Miccheli et al. 2003). These altered metabolic pathways are observable using MRS, as many of the compounds that comprise the NMR-visible neurochemical profile are heavily implicated in energy metabolism.

We report age-related neurochemical modifications in the Fischer rat hippocampus, with the most prominent changes in metabolites implicated in anaerobic energy metabolism, antioxidant capacity, and neuroprotection, as well as numerous macromolecule changes. Previous clinical and preclinical MRS studies of age-related changes have produced mixed findings, particularly for the more commonly reported metabolites such as NAA, tCho, and tCr. Some of the differences between studies may be attributed to reporting metabolite ratios instead of absolute concentrations, different handling of the underlying MM signal, or the study of single sex cohorts (for reviews, see (Haga et al. 2009; Cleeland et al. 2019; Febo and Foster 2016). Below, we present a brief comparison of our findings to previous studies and an interpretation of the biological relevance of the age-related alterations in the neurochemical profile of the Fischer rat.

We detected increased Ins (β=0.363), Glc (β=0.324), Lac (β=0.355), and NAAG (β=0.329) with age, as well as decreased GSH (β=-0.335). Age-related increases in Ins and decreased GSH are generally in good agreement with previous studies (Harris et al. 2014; Zhang et al. 2009; Gruber et al. 2008; Paban, Fauvelle, and Alescio-Lautier 2010; Emir et al. 2011), while reports of changes in Glc, Lac, and NAAG concentrations have been significantly more mixed (Duarte, Do, and Gruetter 2014; Dong and Brewer 2019; Harris et al. 2014; Paban, Fauvelle, and Alescio-Lautier 2010; Miccheli et al. 2003; Małgorzata Marjańska et al. 2017). Despite differences between studies, the neurochemical changes that we report are consistent with the known occurrence of Glc hypometabolism and mitochondrial dysfunction with age, resulting in a shift towards anaerobic energy metabolism, decreased antioxidant capacity, and an increased inflammatory response (Camandola and Mattson 2017; Yin et al. 2016; Miccheli et al. 2003; Godbout and Johnson 2009).

#### Myo-Inositol

Due to the much higher concentration of Ins in astrocytes than in neurons, it has been proposed as a glial cell marker, though increases in Ins do not always correlate with increased gliosis so this interpretation must be taken with caution (Brand, Richter-Landsberg, and Leibfritz 1993; Harris et al. 2014; Best, Stagg, and Dennis 2014). An elevated inflammatory profile has been reported in the aging brain, with astrocytes and microglia displaying a more reactive phenotype, possibly partly in response to increased oxidative damage and stress (Godbout and Johnson 2009)

Ins also plays an important role as a metabolic precursor to the phosphoinositol cycle which is involved in signal transduction and cellular regulation, including Glc metabolism, and is, in fact, derived from D-Glucose-6-phosphate (D-G6P) – the first intermediate formed during glucose catabolism via glycolysis (Bevilacqua and Bizzarri 2018; Zhang et al. 2009). As such, our finding of increased Ins in aged rats could be associated with a number of cell processes affecting neuronal survival and function, including increased glial cell reactivity, overproduction of D-G6P, or decreased activity of the phosphoinositol cycle. Given our other findings of increased Glc (precursor to G6P) and decreased GSH (antioxidant), it is likely that the increase is a result of more than one cellular process.

#### Glucose

Glc hypometabolism and mitochondrial dysfunction resulting in decreased energy production in the form of adenosine triphosphate (ATP) is a known feature of aging (Camandola and Mattson 2017; Yin et al. 2016; Miccheli et al. 2003). Reduced glycolytic activity might explain the increased Glc levels in our aged rats but given our other findings of increased Ins (derived from G6P, the first metabolite in the glycolytic pathway) and Lac (derived from pyruvate, the end product of glycolysis), this hypothesis is unlikely. An alternative possibility given our finding of increased Ins is that increased phosphoinositol cycle signalling molecules are being produced, which participate in the Akt pathway regulating Glc uptake and storage (Bevilacqua and Bizzarri 2018; Best, Stagg, and Dennis 2014). Ins and other inositol derivatives also enhance translocation of the Glc transporter, GLUT-4, through the cell membrane independently of Glc uptake, potentially resulting in altered Glc levels (Bridges and Saltiel 2015; Funaki, DiFransico, and Janmey 2006). Considering the challenges in reliably quantifying Glc with ^1^H MRS, and despite our efforts to ensure good spectral quality, our finding of increased Glc with age and subsequent hypotheses are somewhat open to interpretation and require more investigation.

#### Lactate

Lac is an essential intermediate of Glc metabolism. The glycolytic pathway produces two pyruvate molecules, which then participate in either a) aerobic metabolism via oxidation of pyruvate to acetyl-CoA for entry into the TCA cycle and oxidative phosphorylation within the mitochondria, or b) anaerobic metabolism via reduction of pyruvate to Lac. With age, the pool of nicotinamide adenine dinucleotide (NAD and NAD+) available for oxidative reactions decreases and Glc metabolism shifts from aerobic to anaerobic (Yin et al. 2016; Camandola and Mattson 2017; Dong and Brewer 2019). This shift results in less pyruvate being routed towards the TCA cycle, and more towards reduction into Lac via lactate dehydrogenase (LDH), particularly because LDH activity also increases with age as a result of oxidative stress (Ravera et al. 2019; Dong and Brewer 2019). Increased pyruvate reduction to Lac ensures the cytoplasmic pool of NAD+ is being regenerated, which is necessary for maintaining a high glycolytic rate to produce ATP (Camandola and Mattson 2017; McKenna et al. 2012). As such, our finding of increased Lac fits well with known age-related metabolic changes, as well as our finding of increased Glc and decreased GSH, the latter of which will be discussed below.

#### Glutathione

GSH is an important endogenous antioxidant involved in reducing reactive oxygen species and preventing cellular damage. Production of GSH is coupled with that of ascorbate, another CNS antioxidant, wherein both compounds are reduced by NADP+ as part of the pentose phosphate pathway (Camandola and Mattson 2017; McKenna et al. 2012). Reduced oxidative capacity and impaired mitochondrial respiration is consistent with age-related deficits in antioxidant capacity and therefore the decrease in GSH that we see with age (Emir et al. 2011; McKenna et al. 2012). This finding fits well with that of increased Lac - likely through increased LDH activity triggered by oxidative stress - as an attempt to compensate for impaired mitochondrial energy metabolism. Lastly, increased oxidative damage and stress with reduced antioxidant capacity can trigger glial cell reactivity, which may be reflected in the increased levels of Ins in our aged rats (Godbout and Johnson 2009).

#### N-acetylaspartylglutamate

NAAG is the most abundant neurotransmitter in the CNS, is derived from NAA and Glu, and, like its precursors, is primarily localized in neurons (Benarroch 2008). It acts as a neuromodulatory agent to down-regulate neurotransmitter release via metabotropic Glu receptors, as well as to provide protection to neurons exposed to high Glc conditions (Benarroch 2008; Berent-Spillson et al. 2004). Given our finding of increased Glc, as well as this paper’s description of generally detrimental age-related processes in the brain, the role of NAAG as a compensatory neuroprotective agent and its subsequent increase with age is a reasonable inference.

Other frequently reported metabolites do not appear to change with age in the Fischer rat brain, including tCr, NAA, Glu, Gln, and Asp, though our data do show a trend towards decreased PCr, PCr/Cr, Asp, and Asp/Glu. The trending decrease in PCr/Cr that we found has been reported by others ((Harris et al. 2014; Duarte, Do, and Gruetter 2014; Miccheli et al. 2003) and is proposed to be due to increased PCr to Cr conversion, and thus phosphate donation, in an attempt to maintain cellular ATP supply (McKenna et al. 2012; Best, Stagg, and Dennis 2014). This is consistent with our other findings suggesting energy impairment in the aerobic metabolism pathways. These results also support the argument that significant care should be taken when reporting metabolite ratios to total creatine (Zhang et al. 2009; Duarte et al. 2012; Haga et al. 2009).

NAA, Glu, Gln, and Asp are all metabolites with well-defined roles in energy metabolism and neurotransmission and have previously been shown to be altered with age; Asp and Glu are frequently reported to decrease, Gln to increase, and NAA to either decrease or remain unchanged in older populations relative to young (Małgorzata Marjańska et al. 2017; Hädel et al. 2013; Duarte, Do, and Gruetter 2014; Harris et al. 2014; Zahr et al. 2013; Miccheli et al. 2003). All of these metabolites feed into or are downstream products of the TCA cycle; a transaminase reaction in neuronal mitochondria converts Glu and oxaloacetate into Asp and alpha-ketoglutarate, a TCA-cycle intermediate, while aspartate is used for NAA synthesis. As such, changes in these metabolites are typically interpreted as altered TCA cycle activity and therefore impaired mitochondrial bioenergetics (Dong and Brewer 2019; Yin et al. 2016; Benarroch 2008). Our findings of unaltered tCr, NAA, Glu, and Gln with age requires further investigation, though an alternative explanation that we will discuss in the following section is the frequent study of single sex-cohorts whereas our study examined both males and females.

### 4.3 Characterization of changes in individual macromolecule resonances with age

In 1993 and 1994, Behar et al. characterized the “broad background” signal of MRS spectra as high molecular weight cytosolic macromolecules between 0.9 and 4.4 ppm and assigned each resonance to protons on specific amino acids (Behar and Ogino 1993; Behar et al. 1994). However, due to the fact that the broad peaks are likely representative of overlapping multiplets from various amino acids within different proteins, the exact composition of each MM resonance is not known (Behar and Ogino 1993; Behar et al. 1994; Považan et al. 2018; Małgorzata Marjańska et al. 2018).

Proper handling of the MM signal is intensely important for accurate metabolite quantification. Behar et al. demonstrated that the MM peak at 3.0 ppm represents approximately 5% of the intensity of creatine, but as much as 40-50% of the GABA signal in healthy brain tissue (Behar and Ogino 1993). Similarly, Hofmann et al. reported that inclusion of a MM spectra as a basis function for quantification resulted in considerably lower concentrations for metabolites such as Gln, Glu, and PE, with differences of 0.6 to 1.5 mmol/kg, while also reducing the estimated error margin for all major metabolites (Hofmann et al. 2002). Methodological differences in handling of the MM signal may therefore play a large role in the differences between studies regarding reports of metabolite changes with age and sex or gender (Hofmann et al. 2001) and may explain some of the discrepancies between our study and those published previously.

We report increased MM signal in five of the nine MMs that we quantified. Increased MM signal with age in humans has previously been described (Hofmann et al. 2001; Małgorzata Marjańska et al. 2018), wherein the greatest MM differences associated with age occurred for the 1.7 and 2.0 ppm MM resonances, and a notable increase was observed at 3.9 ppm. This is in agreement with our results, wherein the largest increase occurred in MM_2.02_ (β=0.423), followed by MM_1.20_ (β=0.380), MM_1.66_ (β=0.357), MM_0.89_ (β=0.331), and MM_3.84_ (β=0.290). Coupling experiments measured in 2D COSY and J-resolved spectra of dialyzed human cerebral cytosol indicate strong coupling between MM peaks near 1.6 and 3.0 ppm, as well as between those near 0.93 and 2.1 ppm (Behar et al. 1994). While we noted increases in MM_0.89_ and MM_2.02_, we did not see a similar increase at MM_2.97_ alongside the increase at MM_1.66_. Marjanksa et al. also noted a lack of apparent connectivity for the 1.66 and 2.97 ppm peaks suggesting that perhaps additional macromolecules other than those proposed by Behar et al. (Behar et al. 1994) are contributing to the increase at 1.66 ppm (Małgorzata Marjańska et al. 2018).

Regarding the origin of increased MM signal with age, increases in MM peaks near 0.9, 1.3, and 2.0 ppm have also been shown in subacute stroke patients (Hwang et al. 1996; Saunders et al. 1997). While increased concentrations at 0.9 and 1.3 ppm have been proposed to be caused by increased free fatty acids, the increase at 2.0 ppm is thought to be more likely due to increased visibility of cytosolic proteins after cell death (Saunders et al. 1997). As increased cell death is associated with severe imbalance of glycolysis and the pentose phosphate pathway (Camandola and Mattson 2017), which our findings of increased Lac and decreased GSH support, increased cell death, and thus an increase in cytosolic proteins, is a possible explanation for the general increase in MMs with age that we report. If this were the case, we might expect to also see decreased NAA and/or Glu, which is at least visible (though not statistically significant) in our male-only data.

A related interpretation of the MM signal increase is that, upon cell death, increased cytosolic proteins are being broken down into their subsequent amino acids and feeding into the TCA cycle at various points. This is particularly relevant as our data suggest no major alterations in the TCA cycle metabolites, despite hypothetically decreased pyruvate input due to its increased conversion to Lac. The carbon skeletons of all 20 fundamental amino acids can be broken down and funnelled into only seven molecules, all of which are part of the TCA cycle: pyruvate, acetyl-CoA, alpha-KG, succinyl CoA, fumarate, and oxaloacetate ((Turner 1986; Ulrich et al. 2008). For example, proline and arginine contribute to the MM_2.02_ and MM_1.67_ signal respectively, and their carbon skeletons are fated for use in glutamate, and therefore alpha-KG. Leucine and Isoleucine likely comprise part of the MM_0.89_ signal and also feed into synthesis of Acetyl-CoA. Finally, threonine, cysteine, serine, and glycine all comprise MM signals shown to change in our study, and all contribute to pyruvate synthesis, possibly explaining the ability for increased Lac concentration from pyruvate conversion while maintaining a relatively constant concentration of downstream TCA metabolites.

Hypotheses aside, due to the complexity of each MM signal, it is difficult to determine exactly what proteins within the MM signal are changing with age. As such, the global pattern of changes in MM peaks may be more useful as overall biomarkers of health or pathology as opposed to being indicative of a particular mechanism.

### 4.4 Presence of sex-specific effects in Fisher rat hippocampus

To date, very little work has been done on the characterization of sex differences in the neurochemical profile with age, despite age-related neurodegenerative (and other) disorders such as Alzheimer’s disease differring in prevalence and symptom presentation between sexes (Wickens, Bangasser, and Briand 2018; Komoroski et al. 1999; Mazure and Swendsen 2016). In a longitudinal study of C57BL6 mice, sex differences were identified for many metabolites, including ones in which we also report effects of sex (i.e. Glc and tCr), though the direction of these differences was not noted by the authors (Duarte, Do, and Gruetter 2014). In humans, Hadel et al. described higher hippocampal total creatine in females and lower glutamate in males, and noted a steeper decrease in glutamate with age in females (Hädel et al. 2013). We were not powered to assess the significance of age by sex interactions, but our results collapsed across age do indicate higher total creatine in females and a possible age by sex interaction in Glu concentration.

The importance of studying both sexes is highlighted by a direct comparison of our findings for NAAG, NAA, Asp, and Glu to those reported by Harris et al., who studied aging Fischer rats but only used males. If we look only at our male data, there is a visible trend toward decreased Glu and NAA with age, as well as decreased Asp. This male-only data not only replicates findings by Harris et al. (Harris et al. 2014), but it also explains the increase in NAAG that we report, as NAA, Glu, and Asp all contribute to NAAG synthesis. These trends with age were not visible in the female rats we studied, and in fact, are supported by the trending main effect of sex for NAAG, while the sex effect for Glu and NAA was trending at the p-value level only.

Several of the main effects of sex in our study come from the individual MM resonances at 1.20, 1.39, 2.02, and 2.97 ppm, wherein all but MM_1.20_ were increased in females relative to males. To our knowledge, only a single study by Hofmann et al. has attempted to elucidate sex-specific patterns in the MM signal and they reported no differences between genders (Hofmann et al. 2001). However, even the authors state that the likelihood of finding differences due to gender was low as none were observed for the more easily quantifiable metabolite peaks in this particular study. Additionally, given that the study by Hoffmann et al was conducted at 1.5T, it is likely that the MM peaks were simply not well-resolved enough to detect differences due to sex, and that at higher field strengths sex-specific patterns could emerge, which is what we report.

For MRS studies in general, it is possible that due to different handling of the MM signal, sex- (or age) specific changes in MM_2.02_ and MM_2.97_ peak areas were previously partially attributed to overlapping metabolites such as Glu, Gln, GABA, NAA, NAAG, PCr, or Cr. This has certainly been the argument for some of the differences between studies regarding changes with age in various metabolites such as NAA (Hofmann et al. 2001). Given the general lack of understanding as to the defined role of MMs in aging and pathology, it is difficult to conclude what these effects of sex might mean, but seeing as metabolites can show sex-specific patterns, it is not unreasonable for MMs to do the same, either as a compensatory mechanism, or in response to certain metabolic changes.

Of note, Glc, MM_1.20_, and MM_2.02_ were sensitive to changes with age and differences between sexes, further supporting the necessity of studying both sexes, as well as the hypothesis that parameterization of the MM signal into individual resonances may provide additional biomarkers for detecting pathological changes in brain metabolism.

Regarding the origin of the neurochemical differences between males and females that we report, the current methodological approach does not provide adequate information to define the underlying mechanism(s). However, at the neuroendocrine level, the main difference between the sexes after puberty is the diverging concentration of gonadal hormones estrogen and testosterone. This distinction becomes particularly important during the aging process; estrogens (estradiol, estrone, and estriol) are available to male brains throughout their lifespan via aromatization of testosterone, whereas they are unavailable in the brains of post-menopausal women not using estrogen replacement (Rasgon et al. 2001; Gillies and McArthur 2010). Estrogen receptors are present in the rat and human brain, including the hippocampus, and influence processes including neurogenesis, apoptosis, synaptogenesis, neuroinflammation, establishment of neurochemical pathways, and neurotransmitter control (Gillies and McArthur 2010; Duarte-Guterman et al. 2015; Yin et al. 2015). A pilot study by Rasgon et al. indicated that estrogen replacement therapy may protect against regional cerebral metabolic decline in postmenopausal women (Rasgon et al. 2001), while progesterone has been shown to exert neuroprotective effects including attenuation of oxidative injury from glutamate and glucose-deprivation-induced toxicity (Singh and Su 2013). Most relevant to our work, an elegant study in Sprague-Dawley rats demonstrated that by 16-months all rats were classified as perimenopausal, and that this perimenopausal stage (i.e. reduced estrogen regulation) was associated with altered hippocampal gene expression profiles with roles in insulin signalling, glucose metabolism and mitochondrial function, inflammation, and redox balance (Yin et al. 2015).

Overall, additional work is required to understand sex differences in the neurochemical profile, and future studies should continue to consider both sexes, separately and together, particularly in the context of pathology and treatment. In particular, a reproduction of the detailed gene expression analysis by study by Yin et al. in conjunction with MRS data acquisition throughout the male and female lifespan would be extremely informative.

### 4.5 Age- and sex-dependent change in neurochemistry over three versus four time points

The majority of significant effects seen at four time points were replicated using three time point data, and those that were no longer significant using the smaller dataset were at least trending at the q-value level. The effects of sex remained particularly strong. For more details see **Supplementary Tables 8 and 9**. This additional analysis, in conjunction with the observed power analysis, represents the steps taken towards ensuring our results are robust, despite the decreased sample size towards the end of the study. It is clear that the last timepoint at 20-months constitutes an important piece of the puzzle, both in terms of providing additional power, but also in solidifying age-or sex-related changes in neurochemicals that are trending towards significance in data from the first three timepoints.

### 4.6 Limitations

Despite state-of-the-art protocols and quality control, there are several limitations of this study that need to be considered. First, the number of subjects at our last two timepoints does reduce our overall power. To ensure we were adequately powered for generalization to a larger population, we ran a simulated post-hoc power analysis and present all results, along with their calculated power, for the reader to interpret. We also examined sex effects collapsed across all time points rather than examining age by sex interactions, for which we were underpowered. We recommend that future studies are designed to assess age by sex interactions as they are more informative than examining a main effect of sex.

Second, we chose to use water as a reference without considering the potential up to 5% decrease of water concentration in brain tissue with age, as has been noted by other authors (Chang et al. 1996; Duarte, Do, and Gruetter 2014). As such, it is possible that decreased water content may have resulted in an overestimation of metabolite concentrations towards the later time point. However, a change in water content would drive increases with age similarly across all metabolites. Given that we report some metabolites with significant or trending decreases with age, this suggests that changes in water content are not the dominant driving factor in the changes with age that we observe.

An additional well-known limitation to MRS studies is the possibility of altered relaxation effects with age which may affect metabolite quantification. The measurement of the T2 relaxation time of each neurochemical, including water, is an extremely time-consuming process and therefore not frequently performed, particularly as it is influenced by field strength, age, and brain region (Kreis et al. 2005; Małgorzata Marjańska et al. 2018). As such, while we did not measure the age-specific relaxation constants for all neurochemicals, we did measure T1 and T2 relaxation of water in a separate cohort of 10-month Fischer rats to account for relaxation effects specific to our rat strain, acquisition parameters, and region of interest. We also corrected for non-age associated relaxation effects in metabolites and macromolecules by applying correction factors derived from literature values (for more details, see **Supplementary Table 3**).

Regarding age-related relaxation, we also used a short echo time (TE =11 ms) to minimize T2 relaxation effects. Given an echo time of 11 ms, if the water T2 decreased by 10% in older rats (from 49.13 ms to 44.19 ms) it would decrease T2 relaxation by approximately 2.7%, resulting in metabolite concentration estimates that appear 2.7% higher. Since metabolites such as NAA, Cr, and choline have longer T2s than that of water (Otazo et al. 2006), the effect on the metabolite estimate would be even smaller. As age-specific T2s have not been measured for all neurochemicals at 7T in the rat hippocampus, correcting for age-associated differences in T2 was not possible.

Finally, in comparison to the use of a single-component MM spectrum, the parameterization process of a MM spectrum is lengthy and subject to overfitting during the quantification process if soft constraints are not properly implemented. The complexity of the process can be a deterrent for researchers looking to include individual MMs in their basis set. As such, we have included extensive methods and supplementary material describing our process, in the hopes that other authors will find it helpful for their own studies.

## 5. CONCLUSION

The present study reports, for the first time, the simultaneous absolute quantification of metabolites and individual macromolecules in the aging Fischer rat brain, measured longitudinally using high field ^1^H-MRS. The addition of modelled macromolecule resonances to the standard array of metabolites measured by ^1^H-MRS not only significantly improves metabolite quantification, but also expands the number of potential biomarkers available for detecting pathological changes in brain tissue metabolism. Using this expanded basis set of 31 neurochemicals we identified age- and sex-related changes in tissue chemistry in the hippocampus. The most prominent differences were in metabolites implicated in anaerobic energy metabolism, antioxidant defenses, and neuroprotection, as well as numerous macromolecule changes.

## Supporting information

Supplementary Tables 1 through 9

Supplementary Methods and Equations

## ACKNOWLEDGEMENTS

Sincere thanks are extended to Dana Goerzen and Katrina Cruickshank for their help in editing this manuscript.

## REFERENCES

Bartha, Robert. 2007. “Effect of Signal-to-Noise Ratio and Spectral Linewidth on Metabolite Quantification at 4 T.” NMR in Biomedicine 20 (5): 512–21. https://doi.org/10.1002/nbm.1122.

Behar, K. L., and T. Ogino. 1993. “Characterization of Macromolecule Resonances in the 1H NMR Spectrum of Rat Brain.” Magnetic Resonance in Medicine: Official Journal of the Society of Magnetic Resonance in Medicine / Society of Magnetic Resonance in Medicine 30 (1): 38–44. https://doi.org/10.1002/mrm.1910300107.

Behar, K. L., D. L. Rothman, D. D. Spencer, and O. A. Petroff. 1994. “Analysis of Macromolecule Resonances in 1H NMR Spectra of Human Brain.” Magnetic Resonance in Medicine: Official Journal of the Society of Magnetic Resonance in Medicine / Society of Magnetic Resonance in Medicine 32 (3): 294–302. https://doi.org/10.1002/mrm.1910320304.

Benarroch, E. E. 2008. “N-Acetylaspartate and N-Acetylaspartylglutamate: Neurobiology and Clinical Significance.” Clinical Implications of Neuroscience Research 70 (16): 1353–57. https://doi.org/10.1212/01.wnl.0000311267.63292.6c.

Benjamini, Yoav, and Yosef Hochberg. 1995. “Controlling the False Discovery Rate: A Practical and Powerful Approach to Multiple Testing.” Journal of the Royal Statistical Society. Series B, Statistical Methodology 57 (1): 289–300. http://www.jstor.org/stable/2346101.

Berent-Spillson, Alison, Amanda M. Robinson, David Golovoy, Barbara Slusher, Camilo Rojas, and James W. Russell. 2004. “Protection against Glucose-Induced Neuronal Death by NAAG and GCP II Inhibition Is Regulated by mGluR3.” Journal of Neurochemistry 89 (1): 90–99. https://doi.org/10.1111/j.1471-4159.2003.02321.x.

Best, Jonathan G., Charlotte J. Stagg, and Andrea Dennis. 2014. “Chapter 2.5 - Other Significant Metabolites: Myo-Inositol, GABA, Glutamine, and Lactate.” In Magnetic Resonance Spectroscopy, edited by Charlotte Stagg and Douglas Rothman, 122–38. San Diego: Academic Press. https://doi.org/10.1016/B978-0-12-401688-0.00010-0.

Bettio, Luis E. B., Luckshi Rajendran, and Joana Gil-Mohapel. 2017. “The Effects of Aging in the Hippocampus and Cognitive Decline.” Neuroscience and Biobehavioral Reviews 79 (August): 66–86. https://doi.org/10.1016/j.neubiorev.2017.04.030.

Bevilacqua, Arturo, and Mariano Bizzarri. 2018. “Inositols in Insulin Signaling and Glucose Metabolism.” International Journal of Endocrinology 2018 (November): 1968450. https://doi.org/10.1155/2018/1968450.

Brand, A., C. Richter-Landsberg, and D. Leibfritz. 1993. “Multinuclear NMR Studies on the Energy Metabolism of Glial and Neuronal Cells.” Developmental Neuroscience 15: 289–98. https://doi.org/10.1159/000111347.

Bridges, Dave, and Alan R. Saltiel. 2015. “Phosphoinositides: Key Modulators of Energy Metabolism.” Biochimica et Biophysica Acta 1851 (6): 857–66. https://doi.org/10.1016/j.bbalip.2014.11.008.

Camandola, Simonetta, and Mark P. Mattson. 2017. “Brain Metabolism in Health, Aging, and Neurodegeneration.” The EMBO Journal 36 (11): 1474–92. https://doi.org/10.15252/embj.201695810.

Chang, Linda, Thomas Ernst, Russell E. Poland, and Donald J. Jenden. 1996. “In Vivo Proton Magnetic Resonance Spectroscopy of the Normal Aging Human Brain.” Life Sciences 58 (22): 2049–56. https://doi.org/10.1016/0024-3205(96)00197-x.

Choi, Ji-Kyung, Isabel Carreras, Nur Aytan, Eric Jenkins-Sahlin, Alpaslan Dedeoglu, and Bruce G. Jenkins. 2014. “The Effects of Aging, Housing and Ibuprofen Treatment on Brain Neurochemistry in a Triple Transgene Alzheimer’s Disease Mouse Model Using Magnetic Resonance Spectroscopy and Imaging.” Brain Research 1590 (November): 85–96. https://doi.org/10.1016/j.brainres.2014.09.067.

Cleeland, Carlee, Andrew Pipingas, Andrew Scholey, and David White. 2019. “Neurochemical Changes in the Aging Brain: A Systematic Review.” Neuroscience and Biobehavioral Reviews 98 (March): 306–19. https://doi.org/10.1016/j.neubiorev.2019.01.003.

Craveiro, Mélanie, Virginie Clément-Schatlo, Denis Marino, Rolf Gruetter, and Cristina Cudalbu. 2014. “In Vivo Brain Macromolecule Signals in Healthy and Glioblastoma Mouse Models: 1H Magnetic Resonance Spectroscopy, Post-Processing and Metabolite Quantification at 14.1 T.” Journal of Neurochemistry 129 (5): 806–15. https://doi.org/10.1111/jnc.12673.

Craveiro, Mélanie, Cristina Cudalbu, and Rolf Gruetter. 2012. “Regional Alterations of the Brain Macromolecule Resonances Investigated in the Mouse Brain Using an Improved Method for the Pre-Processing of the Macromolecular Signal.” Proc. Intl. Soc. Mag. Reson. Med. 20: 1748.

Cudalbu, Cristina, Vladimir Mlynárik, and Rolf Gruetter. 2012. “Handling Macromolecule Signals in the Quantification of the Neurochemical Profile.” Journal of Alzheimer’s Disease: JAD 31 Suppl 3: S101–15. https://doi.org/10.3233/JAD-2012-120100.

Dhamala, Elvisha, Ines Abdelkefi, Mavesa Nguyen, T. Jay Hennessy, Hélène Nadeau, and Jamie Near. 2019. “Validation of in Vivo MRS Measures of Metabolite Concentrations in the Human Brain.” NMR in Biomedicine 32 (3): e4058. https://doi.org/10.1002/nbm.4058.

Dong, Yue, and Gregory J. Brewer. 2019. “Global Metabolic Shifts in Age and Alzheimer’s Disease Mouse Brains Pivot at NAD+/NADH Redox Sites.” Journal of Alzheimer’s Disease: JAD 71 (1): 119–40. https://doi.org/10.3233/JAD-190408.

Duarte-Guterman, Paula, Shunya Yagi, Carmen Chow, and Liisa A. M. Galea. 2015. “Hippocampal Learning, Memory, and Neurogenesis: Effects of Sex and Estrogens across the Lifespan in Adults.” Hormones and Behavior 74 (August): 37–52. https://doi.org/10.1016/j.yhbeh.2015.05.024.

Duarte, João M. N., Kim Q. Do, and Rolf Gruetter. 2014. “Longitudinal Neurochemical Modifications in the Aging Mouse Brain Measured in Vivo by 1H Magnetic Resonance Spectroscopy.” Neurobiology of Aging 35 (7): 1660–68. https://doi.org/10.1016/j.neurobiolaging.2014.01.135.

Duarte, João M. N., Hongxia Lei, Vladimír Mlynárik, and Rolf Gruetter. 2012. “The Neurochemical Profile Quantified by in Vivo 1H NMR Spectroscopy.” NeuroImage 61 (2): 342–62. https://doi.org/10.1016/j.neuroimage.2011.12.038.

Emir, Uzay E., Susan Raatz, Susan McPherson, James S. Hodges, Carolyn Torkelson, Pierre Tawfik, Tonya White, and Melissa Terpstra. 2011. “Noninvasive Quantification of Ascorbate and Glutathione Concentration in the Elderly Human Brain.” NMR in Biomedicine 24 (7): 888–94. https://doi.org/10.1002/nbm.1646.

Ernst, T., R. Kreis, and B. D. Ross. 1993. “Absolute Quantitation of Water and Metabolites in the Human Brain. I. Compartments and Water.” Journal of Magnetic Resonance. Series B 102 (1): 1–8. https://doi.org/10.1006/jmrb.1993.1055.

Febo, Marcelo, and Thomas C. Foster. 2016. “Preclinical Magnetic Resonance Imaging and Spectroscopy Studies of Memory, Aging, and Cognitive Decline.” Frontiers in Aging Neuroscience 8 (June): 158. https://doi.org/10.3389/fnagi.2016.00158.

Funaki, Makoto, Lesley DiFransico, and Paul A. Janmey. 2006. “PI 4,5-P2 Stimulates Glucose Transport Activity of GLUT4 in the Plasma Membrane of 3T3-L1 Adipocytes.” Biochimica et Biophysica Acta 1763 (8): 889–99. https://doi.org/10.1016/j.bbamcr.2006.05.012.

Gillies, Glenda E., and Simon McArthur. 2010. “Estrogen Actions in the Brain and the Basis for Differential Action in Men and Women: A Case for Sex-Specific Medicines.” Pharmacological Reviews 62 (2): 155–98. https://doi.org/10.1124/pr.109.002071.

Godbout, Jonathan P., and Rodney W. Johnson. 2009. “Age and Neuroinflammation: A Lifetime of Psychoneuroimmune Consequences.” Immunology and Allergy Clinics of North America 29 (2): 321–37. https://doi.org/10.1016/j.iac.2009.02.007.

Govindaraju, V., K. Young, and A. A. Maudsley. 2000. “Proton NMR Chemical Shifts and Coupling Constants for Brain Metabolites.” NMR in Biomedicine 13 (3): 129–53. https://doi.org/3.0.co;2-v”>10.1002/1099-1492(200005)13:3<129::aid-nbm619>3.0.co;2-v.

Green, Peter, and Catriona J. MacLeod. 2016. “SIMR : An R Package for Power Analysis of Generalized Linear Mixed Models by Simulation.” Edited by Shinichi Nakagawa. Methods in Ecology and Evolution / British Ecological Society 7 (4): 493–98. https://doi.org/10.1111/2041-210X.12504.

Gruber, S., K. Pinker, F. Riederer, M. Chmelik, A. Stadlbauer, M. Bittsansky, V. Mlynarik, et al. 2008. “Metabolic Changes in the Normal Ageing Brain: Consistent Findings from Short and Long Echo Time Spectroscopy.” European Journal of Radiology 68: 320–27. https://doi.org/10.1016/j.ejrad.2007.08.038.

Gruetter, Rolf. 1993. “Automatic, Localized in Vivo Adjustment of All First- and Second-Order Shim Coils.” Journal of Magnetic Resonance in Medicine 29: 804–11. https://doi.org/10.1002/mrm.1910290613.

Hädel, Sven, Christoph Wirth, Michael Rapp, Jürgen Gallinat, and Florian Schubert. 2013. “Effects of Age and Sex on the Concentrations of Glutamate and Glutamine in the Human Brain: Brain Glutamate and Glutamine With Age and Sex.” Journal of Magnetic Resonance Imaging: JMRI 38 (6): 1480–87. https://doi.org/10.1002/jmri.24123.

Haga, Kristin K., Yuet Peng Khor, Andrew Farrall, and Joanna M. Wardlaw. 2009. “A Systematic Review of Brain Metabolite Changes, Measured with 1H Magnetic Resonance Spectroscopy, in Healthy Aging.” Neurobiology of Aging 30 (3): 353–63. https://doi.org/10.1016/j.neurobiolaging.2007.07.005.

Harris, Janna L., Hung-Wen Yeh, Russell H. Swerdlow, In-Young Choi, Phil Lee, and William M. Brooks. 2014. “High-Field Proton Magnetic Resonance Spectroscopy Reveals Metabolic Effects of Normal Brain Aging.” Neurobiology of Aging 35 (7): 1686–94. https://doi.org/10.1016/j.neurobiolaging.2014.01.018.

Hofmann, L., J. Slotboom, C. Boesch, and R. Kreis. 2001. “Characterization of the Macromolecule Baseline in LocalizedH-MR Spectra of Human Brain.” Magnetic Resonance in Medicine: Official Journal of the Society of Magnetic Resonance in Medicine / Society of Magnetic Resonance in Medicine 46: 855–63.

Hofmann, L., J. Slotboom, B. Jung, P. Maloca, C. Boesch, and R. Kreis. 2002. “Quantitative 1H-Magnetic Resonance Spectroscopy of Human Brain: Influence of Composition and Parameterization of the Basis Set in Linear Combination Model-Fitting.” Magnetic Resonance in Medicine: Official Journal of the Society of Magnetic Resonance in Medicine / Society of Magnetic Resonance in Medicine 48 (3): 440–53. https://doi.org/10.1002/mrm.10246.

Hwang, J. H., G. D. Graham, K. L. Behar, J. R. Alger, J. W. Prichard, and D. L. Rothman. 1996. “Short Echo Time Proton Magnetic Resonance Spectroscopic Imaging of Macromolecule and Metabolite Signal Intensities in the Human Brain.” Magnetic Resonance in Medicine: Official Journal of the Society of Magnetic Resonance in Medicine / Society of Magnetic Resonance in Medicine 35 (5): 633–39. https://doi.org/10.1002/mrm.1910350502.

Komoroski, R. A., C. Heimberg, D. Cardwell, and C. N. Karson. 1999. “Effects of Gender and Region on Proton MRS of Normal Human Brain.” Magnetic Resonance Imaging 17 (3): 427–33. https://doi.org/10.1016/s0730-725x(98)00186-6.

Kreis, Roland. 2016. “The Trouble with Quality Filtering Based on Relative Cramér-Rao Lower Bounds.” Magnetic Resonance in Medicine: Official Journal of the Society of Magnetic Resonance in Medicine / Society of Magnetic Resonance in Medicine 75 (1): 15–18. https://doi.org/10.1002/mrm.25568.

Kreis, Roland, Johannes Slotboom, Lucie Hofmann, and Chris Boesch. 2005. “Integrated Data Acquisition and Processing to Determine Metabolite Contents, Relaxation Times, and Macromolecule Baseline in Single Examinations of Individual Subjects.” Magnetic Resonance in Medicine: Official Journal of the Society of Magnetic Resonance in Medicine / Society of Magnetic Resonance in Medicine 54 (4): 761–68. https://doi.org/10.1002/mrm.20673.

Lee, Hyeong Hun, and Hyeonjin Kim. 2017. “Parameterization of Spectral Baseline Directly from Short Echo Time Full Spectra in 1 H-MRS.” Magnetic Resonance in Medicine: Official Journal of the Society of Magnetic Resonance in Medicine / Society of Magnetic Resonance in Medicine 78 (3): 836–47. https://doi.org/10.1002/mrm.26502.

Lopez-Kolkovsky, Alfredo L., Sebastien Mériaux, and Fawzi Boumezbeur. 2016. “Metabolite and Macromolecule T1 and T2 Relaxation Times in the Rat Brain in Vivo at 17.2T.” Magnetic Resonance in Medicine: Official Journal of the Society of Magnetic Resonance in Medicine / Society of Magnetic Resonance in Medicine 75 (2): 503–14. https://doi.org/10.1002/mrm.25602.

Mao, Jintong, T. H. Mareci, and E. R. Andrew. 1988. “Experimental Study of Optimal Selective 180 Radiofrequency Pulses.” Journal of Magnetic Resonance 79 (1): 1–10. https://www.sciencedirect.com/science/article/pii/0022236488903174.

Marjańska, Malgorzata, Dinesh K. Deelchand, James S. Hodges, J. Riley McCarten, Laura S. Hemmy, Andrea Grant, and Melissa Terpstra. 2018. “Altered Macromolecular Pattern and Content in the Aging Human Brain.” NMR in Biomedicine 31 (2). https://doi.org/10.1002/nbm.3865.

Marjańska, Malgorzata, J. Riley McCarten, James Hodges, Laura S. Hemmy, Andrea Grant, Dinesh K. Deelchand, and Melissa Terpstra. 2017. “Region-Specific Aging of the Human Brain as Evidenced by Neurochemical Profiles Measured Noninvasively in the Posterior Cingulate Cortex and the Occipital Lobe Using 1H Magnetic Resonance Spectroscopy at 7 T.” Neuroscience 354 (June): 168–77. https://doi.org/10.1016/j.neuroscience.2017.04.035.

Marjańska, Malgorzata, J. Riley McCarten, James S. Hodges, Laura S. Hemmy, and Melissa Terpstra. 2019. “Distinctive Neurochemistry in Alzheimer’s Disease via 7 T In Vivo Magnetic Resonance Spectroscopy.” Journal of Alzheimer’s Disease: JAD 68 (2): 559–69. https://doi.org/10.3233/JAD-180861.

Marjańska, M., S. D. Weigand, G. Preboske, T. M. Wengenack, R. Chamberlain, G. L. Curran, J. F. Poduslo, et al. 2014. “Treatment Effects in a Transgenic Mouse Model of Alzheimer’s Disease: A Magnetic Resonance Spectroscopy Study after Passive Immunization.” Neuroscience 259 (February): 94–100. https://doi.org/10.1016/j.neuroscience.2013.11.052.

Mazure, Carolyn M., and Joel Swendsen. 2016. “Sex Differences in Alzheimer’s Disease and Other Dementias.” Lancet Neurology. https://doi.org/10.1016/S1474-4422(16)00067-3.

McKenna, Mary C., Gerald A. Dienel, Ursula Sonnewald, Helle S. Waagepetersen, and Arne Schousboe. 2012. “Chapter 11 - Energy Metabolism of the Brain.” In Basic Neurochemistry (Eighth Edition), edited by Scott T. Brady, George J. Siegel, R. Wayne Albers, and Donald L. Price, 200–231. New York: Academic Press. https://doi.org/10.1016/B978-0-12-374947-5.00011-0.

Miccheli, Alfredo, Caterina Puccetti, Giorgio Capuani, Maria Enrica Di Cocco, Luciana Giardino, Laura Calzà, Angelo Battaglia, Leontino Battistin, and Filippo Conti. 2003. “[1-13C]Glucose Entry in Neuronal and Astrocytic Intermediary Metabolism of Aged Rats: A Study of the Effects of Nicergoline Treatment by 13C NMR Spectroscopy.” Brain Research 966 (1): 116–25. https://doi.org/10.1016/S0006-8993(02)04217-8.

Mlynárik, Vladimír, Giulio Gambarota, Hanne Frenkel, and Rolf Gruetter. 2006. “Localized Short-Echo-Time Proton MR Spectroscopy with Full Signal-Intensity Acquisition.” Magnetic Resonance in Medicine: Official Journal of the Society of Magnetic Resonance in Medicine / Society of Magnetic Resonance in Medicine 56 (5): 965–70. https://doi.org/10.1002/mrm.21043.

Morrison, John H., Paul C. Stern, and Laura L. Carstensen. 2000. Age-Related Shifts in Neural Circuit Characteristics and Their Impact on Age-Related Cognitive Impairments. National Academies Press (US). https://www.ncbi.nlm.nih.gov/books/NBK44828/.

Murray, Melissa E., Scott A. Przybelski, Timothy G. Lesnick, Amanda M. Liesinger, Anthony Spychalla, Bing Zhang, Jeffrey L. Gunter, et al. 2014. “Early Alzheimer’s Disease Neuropathology Detected by Proton MR Spectroscopy.” The Journal of Neuroscience: The Official Journal of the Society for Neuroscience 34 (49): 16247–55. https://doi.org/10.1523/JNEUROSCI.2027-14.2014.

Naressi, A., C. Couturier, J. M. Devos, M. Janssen, C. Mangeat, R. de Beer, and D. Graveron-Demilly. 2001. “Java-Based Graphical User Interface for the MRUI Quantitation Package.” Magma 12 (2-3): 141–52. https://doi.org/10.1007/bf02668096.

Nilsen, L. H., T. M. Melø, M. P. Witter, and U. Sonnewald. 2014. “Early Differences in Dorsal Hippocampal Metabolite Levels in Males but Not Females in a Transgenic Rat Model of Alzheimer’s Disease.” Neurochemical Research 39 (2): 305–12. https://doi.org/10.1007/s11064-013-1222-x.

Otazo, Ricardo, Bryon Mueller, Kamil Ugurbil, Lawrence Wald, and Stefan Posse. 2006. “Signal-to-Noise Ratio and Spectral Linewidth Improvements between 1.5 and 7 Tesla in Proton Echo-Planar Spectroscopic Imaging.” Magnetic Resonance in Medicine: Official Journal of the Society of Magnetic Resonance in Medicine / Society of Magnetic Resonance in Medicine 56 (6): 1200–1210. https://doi.org/10.1002/mrm.21067.

Paban, Véronique, Florence Fauvelle, and Béatrice Alescio-Lautier. 2010. “Age-Related Changes in Metabolic Profiles of Rat Hippocampus and Cortices.” The European Journal of Neuroscience 31 (6): 1063–73. https://doi.org/10.1111/j.1460-9568.2010.07126.x.

Pfeuffer, Josef, Ivan Tkač, Stephen W. Provencher, and Rolf Gruetter. 1999. “Toward an in Vivo Neurochemical Profile: Quantification of 18 Metabolites in Short-Echo-TimeH NMR Spectra of the Rat Brain.” Journal of Magnetic Resonance 141: 104–20.

Pijnappel, W. W. F., A. van den Boogaart, R. de Beer, and D. van Ormondt. 1992. “SVD-Based Quantification of Magnetic Resonance Signals.” Journal of Magnetic Resonance (1969). https://doi.org/10.1016/0022-2364(92)90241-x.

Považan, Michal, Bernhard Strasser, Gilbert Hangel, Eva Heckova, Stephan Gruber, Siegfried Trattnig, and Wolfgang Bogner. 2018. “Simultaneous Mapping of Metabolites and Individual Macromolecular Components via Ultra-Short Acquisition Delay 1 H MRSI in the Brain at 7T : Simultaneous Mapping of Metabolites and Macromolecules.” Magnetic Resonance in Medicine: Official Journal of the Society of Magnetic Resonance in Medicine / Society of Magnetic Resonance in Medicine 79 (3): 1231–40. https://doi.org/10.1002/mrm.26778.

Provencher, Stephen. 2019. “LCModel & LCMgui User’s Manual.”

Provencher, S. W. 1993. “Estimation of Metabolite Concentrations from Localized in Vivo Proton NMR Spectra.” Magnetic Resonance in Medicine: Official Journal of the Society of Magnetic Resonance in Medicine / Society of Magnetic Resonance in Medicine 30 (6): 672–79. https://doi.org/10.1002/mrm.1910300604.

Rasgon, N. L., G. W. Small, P. Siddarth, K. Miller, L. M. Ercoli, S. Y. Bookheimer, H. Lavretsky, S. C. Huang, J. R. Barrio, and M. E. Phelps. 2001. “Estrogen Use and Brain Metabolic Change in Older Adults. A Preliminary Report.” Psychiatry Research 107 (1): 11–18. https://doi.org/10.1016/s0925-4927(01)00084-1.

Ravera, Silvia, Marina Podestà, Federica Sabatini, Monica Dagnino, Daniela Cilloni, Samuele Fiorini, Annalisa Barla, and Francesco Frassoni. 2019. “Discrete Changes in Glucose Metabolism Define Aging.” Scientific Reports 9 (1): 10347. https://doi.org/10.1038/s41598-019-46749-w.

Ross, Amy J., and Perminder S. Sachdev. 2004. “Magnetic Resonance Spectroscopy in Cognitive Research.” Brain Research. Brain Research Reviews 44 (2-3): 83–102. https://doi.org/10.1016/j.brainresrev.2003.11.001.

Saunders, D. E., F. A. Howe, A. van den Boogaart, J. R. Griffiths, and M. M. Brown. 1997. “Discrimination of Metabolite from Lipid and Macromolecule Resonances in Cerebral Infarction in Humans Using Short Echo Proton Spectroscopy.” Journal of Magnetic Resonance Imaging: JMRI 7 (6): 1116–21. https://doi.org/10.1002/jmri.1880070626.

Schuff, Norbert, Diane L. Amend, Robert Knowlton, David Norman, George Fein, and Michael W. Weiner. 1999. “Age-Related Metabolite Changes and Volume Loss in the Hippocampus by Magnetic Resonance Spectroscopy and Imaging<.” Neurobiology of Aging 20: 279–85.

Seeger, Uwe, Uwe Klose, Irina Mader, Wolfgang Grodd, and Thomas Nägele. 2003. “Parameterized Evaluation of Macromolecules and Lipids in Proton MR Spectroscopy of Brain Diseases.” Magnetic Resonance in Medicine: Official Journal of the Society of Magnetic Resonance in Medicine / Society of Magnetic Resonance in Medicine 49 (1): 19–28. https://doi.org/10.1002/mrm.10332.

Simpson, Robin, Gabriel A. Devenyi, Peter Jezzard, T. Jay Hennessy, and Jamie Near. 2017. “Advanced Processing and Simulation of MRS Data Using the FID Appliance (FID-A)-An Open Source, MATLAB-Based Toolkit.” Magnetic Resonance in Medicine: Official Journal of the Society of Magnetic Resonance in Medicine / Society of Magnetic Resonance in Medicine 77 (1): 23–33. https://doi.org/10.1002/mrm.26091.

Singh, Meharvan, and Chang Su. 2013. “Progesterone and Neuroprotection.” Hormones and Behavior 63 (2): 284–90. https://doi.org/10.1016/j.yhbeh.2012.06.003.

Snoussi, Karim, Joseph S. Gillen, Alena Horska, Nicolaas A. J. Puts, Subechhya Pradhan, Richard A. E. Edden, and Peter B. Barker. 2015. “Comparison of Brain Gray and White Matter Macromolecule Resonances at 3 and 7 Tesla.” Magnetic Resonance in Medicine: Official Journal of the Society of Magnetic Resonance in Medicine / Society of Magnetic Resonance in Medicine 74 (3): 607–13. https://doi.org/10.1002/mrm.25468.

Stefan, D., F. Di Cesare, A. Andrasescu, E. Popa, A. Lazariev, E. Vescovo, O. Strbak, et al. 2009. “Quantitation of Magnetic Resonance Spectroscopy Signals: The jMRUI Software Package.” Measurement Science & Technology 20 (10): 104035. https://doi.org/10.1088/0957-0233/20/10/104035.

Tkač, I., Z. Starčuk, I. -Y. Choi, and R. Gruetter. 1999. “In VivoH NMR Spectroscopy of Rat Brain at 1 Ms Echo Time.” Magnetic Resonance in Medicine: Official Journal of the Society of Magnetic Resonance in Medicine / Society of Magnetic Resonance in Medicine 41: 649–56.

Turner, A. J. 1986. “Biochemistry and the Central Nervous System (Fifth Edition).” Biochemical Education 14 (1): 46. https://doi.org/10.1016/0307-4412(86)90054-3.

Ulrich, Eldon L., Hideo Akutsu, Jurgen F. Doreleijers, Yoko Harano, Yannis E. Ioannidis, Jundong Lin, Miron Livny, et al. 2008. “BioMagResBank.” Nucleic Acids Research 36 (Database issue): D402–8. https://doi.org/10.1093/nar/gkm957.

Vanhamme, L., van den Boogaart A, and Van Huffel S. 1997. “Improved Method for Accurate and Efficient Quantification of MRS Data with Use of Prior Knowledge.” Journal of Magnetic Resonance 129 (1): 35–43. https://doi.org/10.1006/jmre.1997.1244.

Van Hoesen, G. W., B. T. Hyman, and A. R. Damasio. 1991. “Entorhinal Cortex Pathology in Alzheimer’s Disease.” Hippocampus 1 (1): 1–8. https://doi.org/10.1002/hipo.450010102.

Wickens, Megan M., Debra A. Bangasser, and Lisa A. Briand. 2018. “Sex Differences in Psychiatric Disease: A Focus on the Glutamate System.” Frontiers in Molecular Neuroscience 11 (June): 197. https://doi.org/10.3389/fnmol.2018.00197.

Yin, Fei, Harsh Sancheti, Ishan Patil, and Enrique Cadenas. 2016. “Energy Metabolism and Inflammation in Brain Aging and Alzheimer’s Disease.” Free Radical Biology & Medicine 100 (November): 108–22. https://doi.org/10.1016/j.freeradbiomed.2016.04.200.

Yin, Fei, Jia Yao, Harsh Sancheti, Tao Feng, Roberto C. Melcangi, Todd E. Morgan, Caleb E. Finch, et al. 2015. “The Perimenopausal Aging Transition in the Female Rat Brain: Decline in Bioenergetic Systems and Synaptic Plasticity.” Neurobiology of Aging 36 (7): 2282–95. https://doi.org/10.1016/j.neurobiolaging.2015.03.013.

Zahr, Natalie M., Dirk Mayer, Torsten Rohlfing, Sandra Chanraud, Meng Gu, Edith V. Sullivan, and Adolf Pfefferbaum. 2013. “In Vivo Glutamate Measured with Magnetic Resonance Spectroscopy: Behavioral Correlates in Aging.” Neurobiology of Aging 34 (4): 1265–76. https://doi.org/10.1016/j.neurobiolaging.2012.09.014.

Zhang, Xianrong, Huilang Liu, Junfang Wu, Xu Zhang, Maili Liu, and Yong Wang. 2009. “Metabonomic Alterations in Hippocampus, Temporal and Prefrontal Cortex with Age in Rats.” Neurochemistry International 54 (8): 481–87. https://doi.org/10.1016/j.neuint.2009.02.004.

