## Supplementary Methods and Equations for "Longitudinal Quantification of Metabolites and Macromolecules Reveals Age- and Sex-Related Changes in the Healthy Fischer 344 Rat Brain"

**Quality Control**

Each spectrum was visually inspected for quality before and after post-processing with FID-A. The full-width half-max (FWHM) of the water unsuppressed spectrum and the SNR of the NAA peak in the water suppressed spectrum was extracted from each spectrum using an in-house MATLAB (R2012a) script (below) in conjunction with the FID-A toolkit (relevant commands: op_getSNR, and op_getLW). Average linewidth and SNR for each timepoint are shown in **Supplementary Table 5,** alongside values for estimated spectral FWHM and SNR for water suppressed data provided by LCModel. Quality control resulted in the discarding of a single spectrum at the second time point due to SNR lower than 25, reducing the number of scans from 30 to 29.

**Processing of macromolecule spectra**

MM scans were obtained in 8 Fischer rats from a separate cohort at 10-months of age (313.8 ± 11.7 days). First, $^1$H-MRS scans were acquired after FASTMAP shimming and a water reference scan. Next, “metabolite-suppressed” spectra were acquired using PRESS localization preceded by an IR pulse with an inversion time (TI) of 800 ms. In each rat, the metabolite-suppressed IR scan was acquired immediately after the $^1$H-MRS scan in order to avoid re-shimming the instrument.

For each rat, water unsuppressed data, water suppressed data, and macromolecule data were imported into the FID-A processing toolbox (version 1.0) in MATLAB (R2012a). Within each dataset, transients were averaged and a zero-order phase shift correction was applied, using the phase of the NAA peak within the metabolite spectra. The NAA peak was also used to determine the frequency shift, and both the determined frequency and phase shifts were then applied to the water suppressed and macromolecule data. The eight macromolecule spectra were then aligned, summed, and scaled to generate an average metabolite-nulled spectra, for parameterization.

**Determination of Water and Metabolite Relaxation Constants for Absolute Quantification**

T1 and T2 water relaxation constants for use in absolute quantification were acquired in a subset of Fischer rats (n=7) at 10-months of age. The T2 relaxation constant of water was determined by acquiring a series of non-localized water spectra in the same region of interest described above, using the PRESS sequence (TR = 5000 ms, 8 averages) without water suppression and with varying echo times (12, 20, 30, 40, 50, 70, 90, 120, 150, 200, 300, and 500 ms). Mono-exponential fitting of the T2 curves provided an estimated water T2 of 49.13 ms. To determine the T1 relaxation of water, a series of non-localized water spectra were acquired using the PRESS sequence (TR = 5000ms, TE=12 ms, 8 averages) preceded by a 4th order hyperbolic secant adiabatic full passage (AFP) inversion pulse (pulse duration = 1.0ms, bandwidth = 6982.0 Hz). A series of inversion times (TI) (25, 94, 261, 273, 660, 910, 1160, 1409, 1660, 2160, 3160, and 4160 ms) was acquired, with an additional scan performed without the inversion pulse to determine maximum signal intensity. The average maximum signal intensity of water at different inversion times (S(TI)) across the 7 rats were fitted versus their corresponding TIs using the equation below (Dehghani et al. 2020) resulting in an estimated water T1 of 1491ms.

**Supplementary Equation 1:**

$$S(TI) = S(TI_{off})\cdot(1 - (1 + \alpha)\cdot exp^{-TI/T1} + \alpha\cdot exp^{-TR/T1}) / (1 - exp^{-TR/T1}),$$

Where α represents the factor for the flip angle of the inversion pulse, and $S(TI_{off})$ represents the signal intensity at TE of 12 ms without inversion.

To approximate the T1 and T2 constants of individual metabolites at 7T, we fit a simple linear model (y = a*x + b) to values obtained by de Graaf et al. for tCho, tCr, Glx, NAA, and macromolecules, at 4 and 9.4T (de Graaf et al. 2006), For relaxation constants in metabolites not specifically measured by de Graaf, we used the average across projected metabolite relaxation constants.

The following equation was used to account for the T1 and T2 relaxation constants of water, metabolites and macromolecules. **Supplementary Equation 2**, modified from Dhamala et al., (Dhamala et al. 2019), also includes a correction factor to account for the fact that our voxel contained primarily grey matter (with an NMR-visible water concentration of 43300 mM) as opposed to the default for white matter used by LCModel (35880 mM). All relaxation constants and the subsequent correction factors applied, are summarized in **Supplementary Table 3**.

**Supplementary Equation 2:**

$$metabolite.abs = (signal_{met} / signal_{H2O}) \times (WCONC_{GMH2O}) \times [ (exp^{-TE/T2H2O}\cdot(1 - exp^{-TR/T1H2O})) / (exp^{-TE/T2met}\cdot(1 - exp^{-TR/T1met})) ],$$

Where *metabolite.abs* is the absolute concentration of a given metabolite.

$signal_{met}/signal_{H2O}$ is the ratio of metabolite signal to water signal, as determined using LCmodel. This value is returned by LCModel when the parameters WCONC, ATTH2O, and ATTMET are all set to 1, and water scaling is on.

$WCONC_{GMH2O}$ is the LCModel parameter specifying the tissue water concentration in grey matter (43300 mM) (Ernst, Kreis, and Ross 1993).

*TE* is the echo time of the experiment (TE = 11.12 ms)

*TR* is the repetition time of the experiment (TR = 3000 ms)

$T2_{H2O}$ is the measured water T2 relaxation time at 7T (49.13 ms)

$T2_{met}$ is the projected metabolite T2 relaxation time at 7T (**Supplementary Table 3**)

$T1_{H2O}$ is the measured water T1 relaxation time at 7T (1491 ms)

$T1_{met}$ is projected metabolite T2 relaxation time at 7T (**Supplementary Table 3**)

#### Retrospective Power Analysis

Retrospective power analyses to determine if the absence of an effect was due to lack of power are typically not advised, as there is a direct relationship between observed power and p-values (Hoenig and Heisey, n.d.). However, we are attempting to do the opposite and demonstrate that the effects we do see in our relatively small sample size are, in fact, generalizable to a larger population, i.e. given the variability within our dataset, if the data is simulated *x* number of times, how often is the effect size of interest statistically significant. This is particularly important because our dataset has a large reduction in subjects at the last two timepoints. As such, power calculations were performed in SIMR using Monte Carlo simulations (n=1000) and produced a calculated power and 95% confidence interval for the fixed effects of sex (collapsed across timepoints) and age for each metabolite. Traditionally 80% power is considered adequate, though as this is a somewhat arbitrary cut-off (Bacchetti 2010) we present all results, including the observed power and effect size (standardized beta) for each metabolite, for the reader to interpret as they wish.

### REFERENCES:


Bacchetti, Peter. 2010. “Current Sample Size Conventions: Flaws, Harms, and Alternatives.” *BMC Medicine* 8 (March): 17.

Dehghani, Masoumeh, Kim Q. Do, Pierre Magistretti, and Lijing Xin. 2020. “Lactate Measurement by Neurochemical Profiling in the Dorsolateral Prefrontal Cortex at 7T: Accuracy, Precision, and Relaxation Times.” *Magnetic Resonance in Medicine: Official Journal of the Society of Magnetic Resonance in Medicine / Society of Magnetic Resonance in Medicine* 83 (6): 1895–1908.

Dhamala, Elvisha, Ines Abdelkefi, Mavesa Nguyen, T. Jay Hennessy, Hélène Nadeau, and Jamie Near. 2019. “Validation of in Vivo MRS Measures of Metabolite Concentrations in the Human Brain.” *NMR in Biomedicine* 32 (3): e4058.

Ernst, T., R. Kreis, and B. D. Ross. 1993. “Absolute Quantitation of Water and Metabolites in the Human Brain. I. Compartments and Water.” *Journal of Magnetic Resonance. Series B* 102 (1): 1–8.

Graaf, Robin A. de, Peter B. Brown, Scott McIntyre, Terence W. Nixon, Kevin L. Behar, and Douglas L. Rothman. 2006. “High Magnetic Field Water and Metabolite Proton T1 and T2 Relaxation in Rat Brain in Vivo.” *Magnetic Resonance in Medicine: Official Journal of the Society of Magnetic Resonance in Medicine / Society of Magnetic Resonance in Medicine* 56 (2): 386–94.

Hoenig, John M., and Dennis M. Heisey. n.d. “The Abuse of Power: The Pervasive Fallacy of Power Calculations for Data Analysis.”
